# Fat Body Phospholipid State Dictates Hunger Driven Feeding Behavior

**DOI:** 10.1101/2021.12.16.472854

**Authors:** Kevin P. Kelly, Mroj Alassaf, Camille E. Sullivan, Ava E. Brent, Zachary H. Goldberg, Michelle E. Poling, Julien Dubrulle, Akhila Rajan

**Author notes:** Equal Contribution (Kevin P. Kelly, Mroj Alassaf).

## Abstract

Diet-induced obesity (DIO) leads to dysfunctional feeding behavior. However, the precise molecular nodes underlying diet-induced dysregulation of satiety sensing and feeding motivation are poorly understood. The fruit fly is a simple genetic model system yet displays significant evolutionary conservation to mammalian nutrient sensing and energy balance. Using a longitudinal high sugar regime in *Drosophila*, we sought to address how lipid alteration in fat cells alters feeding behavior. We find that prolonged exposure to HSD degrades the hunger-driven feeding (HDF) response. Lipidomics analysis reveals that longitudinal exposure to HSD significantly alters whole body phospholipid profiles. By performing a systematic screen for phospholipid enzymes, we identify Pect as a critical regulator of hunger-driven feeding. Pect is a rate-limiting enzyme in the phosphatidylethanolamine (PE) biosynthesis pathway and the fly ortholog of human PCYT2. We show that disrupting Pect only in the fat body causes insulin-resistant phenotypes and a loss of hunger-driven feeding. Excitingly, we find that overexpression of Pect restores HSD-induced loss of hunger-driven feeding response. Strikingly human studies have noted a correlation between PCYT2/Pect levels and clinical obesity. Now, our unbiased studies in *Drosophila* provide specific genetic evidence for Pect in maintaining nutrient sensing during DIO. Our study provides novel insights into the role of phospholipids in interorgan communication of nutrient status.

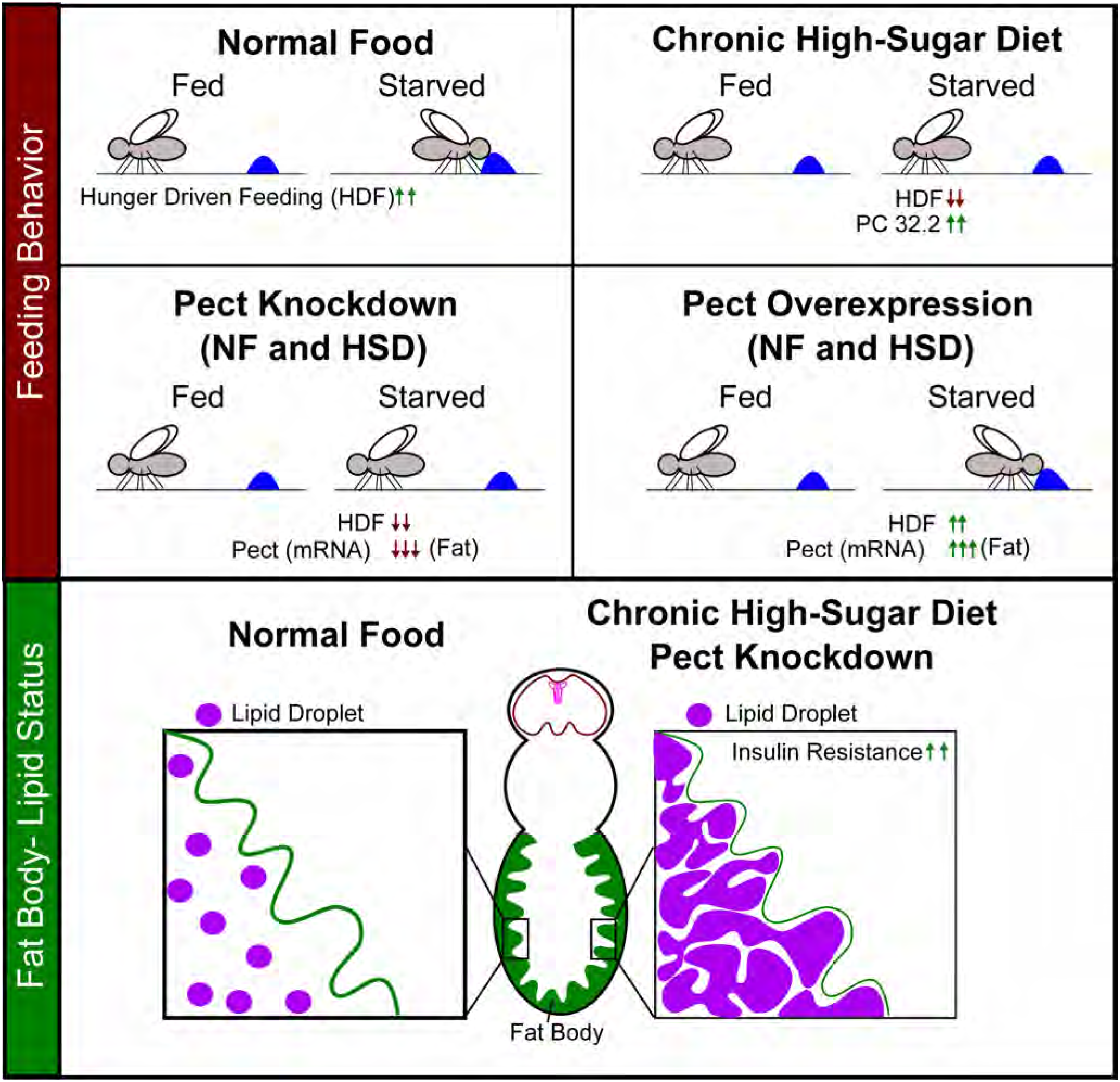

## Introduction

The growing prevalence of diet-induced obesity (DIO) and its associated comorbidities such as type 2 diabetes mellitus place a significant burden on the healthcare system^1^. Hunger and feeding behavior are coupled to facilitate adequate nutritional intake^2^. In DIO, disrupted hunger perception leads to dysfunctional feeding behavior^3^. While the neural circuits regulating hunger are becoming well understood^4^, our understanding of how DIO disrupts adipose tissue function and impact hunger and feeding behavior is only now emerging^5^. In addition to its role in lipid storage, adipose tissue serves as a major endocrine and secretory organ^6,7^. Adipocyte-secreted factors impinge on several organs, including the brain to regulate systemic metabolism and feeding behavior^8–11^. Therefore, understanding the mechanisms underlying diet-induced adipocyte dysfunction, and how it impacts feeding behavior, may provide a potential therapeutic target against obesity. Adipocyte-secreted factors impinge on several organs, including the brain to regulate systemic metabolism and feeding behavior^8–11^. Therefore, understanding the mechanisms underlying diet-induced adipocyte dysfunction, and how it impacts feeding behavior, may provide a potential therapeutic target against obesity.

The effects of several base components of food, such as triglycerides, on feeding behavior have been extensively studied^12^. However, less is known about the effects of phospholipids. Phospholipids comprise the lipid bilayer of the plasma membrane and anchor integral membrane proteins including ion channels and receptors. They are essential components of cellular organelles, lipoproteins, and secretory vesicles^13^. Changes to phospholipid composition can alter the permissibility of the cellular membrane and disrupt intra- and intercellular signaling^13–15^. Numerous clinical studies point to a link between phospholipid composition and obesity^16–18^. For example, insulin resistance, a hallmark of obesity-induced type 2 diabetes, is strongly associated with alterations in phospholipid composition^17^. Additionally, key phospholipid biosynthesis enzymes are correlated with obesity in human GWAS studies^16^. While there is a strong correlation between obesity and changes to the lipidome, a causative relationship is yet to be elucidated. Given that DIO can impact phospholipid composition and as a result disrupt cell signaling, defects in interorgan communication are possible. Despite these intriguing possibilities, no functional genetic studies have systematically examined the effects of phospholipid dysregulation on feeding behavior.

Phosphatidylethanolamine (PE) is among the second most abundant phospholipid and is essential in membrane fission/fusion events^19–21^. PE is synthesized through two main pathways that occur in the ER and the mitochondria^22^. Phosphatidylethanolamine cytidylyltransferase (Pcyt/Pect) is the rate-limiting enzyme of the ER-mediated PE biosynthesis pathway^23^. Pcyt/Pect activity has been shown to impact systemic metabolism^24,25^. For example, Pyct/Pect deficiency in Pcyt/Pect^+/-^ mice causes a reduction in PE levels which leads to obesity and insulin resistance^26^. Additionally, human studies found that obese individuals with insulin resistance have decreased Pcyt/Pect expression levels^27^. Interestingly, chronic exposure to a high-fat diet causes upregulation of Pcyt/Pect that is associated with increased weight gain and insulin resistance. An analysis of the adipose tissues of mice fed a HFD revealed that Pcyt2 levels were higher in high weight gainers compared to low weight gainers^28^. These findings suggest that dysregulation in PE homeostasis is a common underlying feature of obesity and metabolic disorders, which suggests that Pcyt/Pect may be a vital regulator of systemic metabolism. What remains largely unknown is whether Pcyt/Pect activity and the resultant changes in PE levels can affect feeding behavior.

The components of DIO and insulin resistance are highly evolutionarily conserved in both flies and mammals^29–33^. *Drosophila* is an established metabolic model organism that has been shown to mimic the feeding behavior of humans in response to high palatable foods^34–38^. In contrast to rodents, *Drosophila* do not exhibit hoarding behavior, which allows for accurate measurement of food intake. Additionally, given the short lifespan of flies, feeding behavior can be monitored at all stages of the fly’s adult life, which can provide temporal resolution of behavioral changes under DIO^39–43^. Studies on *Drosophila* feeding behavior and appetite proved successful in identifying key neurons and receptors involved^44–46^. What remains unclear is how peripheral organs communicate with the brain and how this communication is dysregulated in obesogenic diets. Similar to humans, chronic overconsumption of a high sugar diet (HSD) results in insulin resistance, obesity, and metabolic imbalance in flies^47^. Thus, using a chronic HSD feeding regime in adult flies may allow for the discovery of specific mechanisms relevant to human biology.

In this study, we assess the effects of chronic HSD consumption on the hunger-driven feeding (HDF) behavior of flies across a 28-day time window. We note that flies display a significant increase in triglyceride store on HSD exposure and maintain their ability to mobilize fat stores on starvation despite prolonged HSD exposure. But we find that while flies fed a normal diet (NF) display increased feeding upon starvation throughout this 4-week period, HSD-fed flies lose this HDF-response after 2-weeks. We reveal that changes in phospholipid concentrations in HSD-fed flies occur at the time of HDF loss. We further show that genetic disruption of the key PE biosynthesis enzyme Pect in the fat body, the fly’s adipose tissue, results in the loss of HDF. Significantly, Pect overexpression in the fat body is sufficient to protect flies from HSD-induced loss in HDF. Combined, our data suggest that phospholipid homeostasis in adipose tissue is a critical component to the maintenance of hunger-driven feeding behavior.

## Results

### Hunger Driven Feeding in Flies is lost under High Sugar Diet

The inability to sense hunger underlies a multitude of eating disorders, including obesity^48^. Yet, the cellular and molecular mechanisms governing the breakdown of the hunger-sensing system in DIO are unknown. To address this, we developed a *Drosophila* model of HSD-induced Obesity. We first sought to determine if flies exhibit hunger-driven feeding behavior and whether this behavior changes in response to HSD treatment. We used a Fly Liquid Interaction Counter (FLIC)^39^ to monitor fly feeding activity overtime on different diets (Figure 1A). In brief, flies were housed in vials containing NF or HSD until16 hours prior to feeding behavior measurement, after which half of the flies were moved to a 0% sucrose/agar media to induce starvation (Hereby referred to as Stv condition). Flies were then monitored for feeding behavior on FLIC for three hours (10 am-1 pm local time). When placed under 16 hours of starvation media, wild type (w1118) flies showed increased feeding compared to flies allowed to eat *ad libitum* (Figure 1B and Figure S1), which is consistent with a HDF response seen in vertebrates^49,50^. However, flies fed a HSD (30% more sucrose than NF) showed a progressive loss of the HDF response to starvation starting at day 14, suggesting a breakdown in the hunger-sensing system (Figure 1C and Figure S1). Notably, flies on NF and HSD showed similar *ad libitum* feeding behavior excluding the possibility that the loss in HDF is due to increased baseline feeding on HSD (Figure 1D).

**Figure 1.**
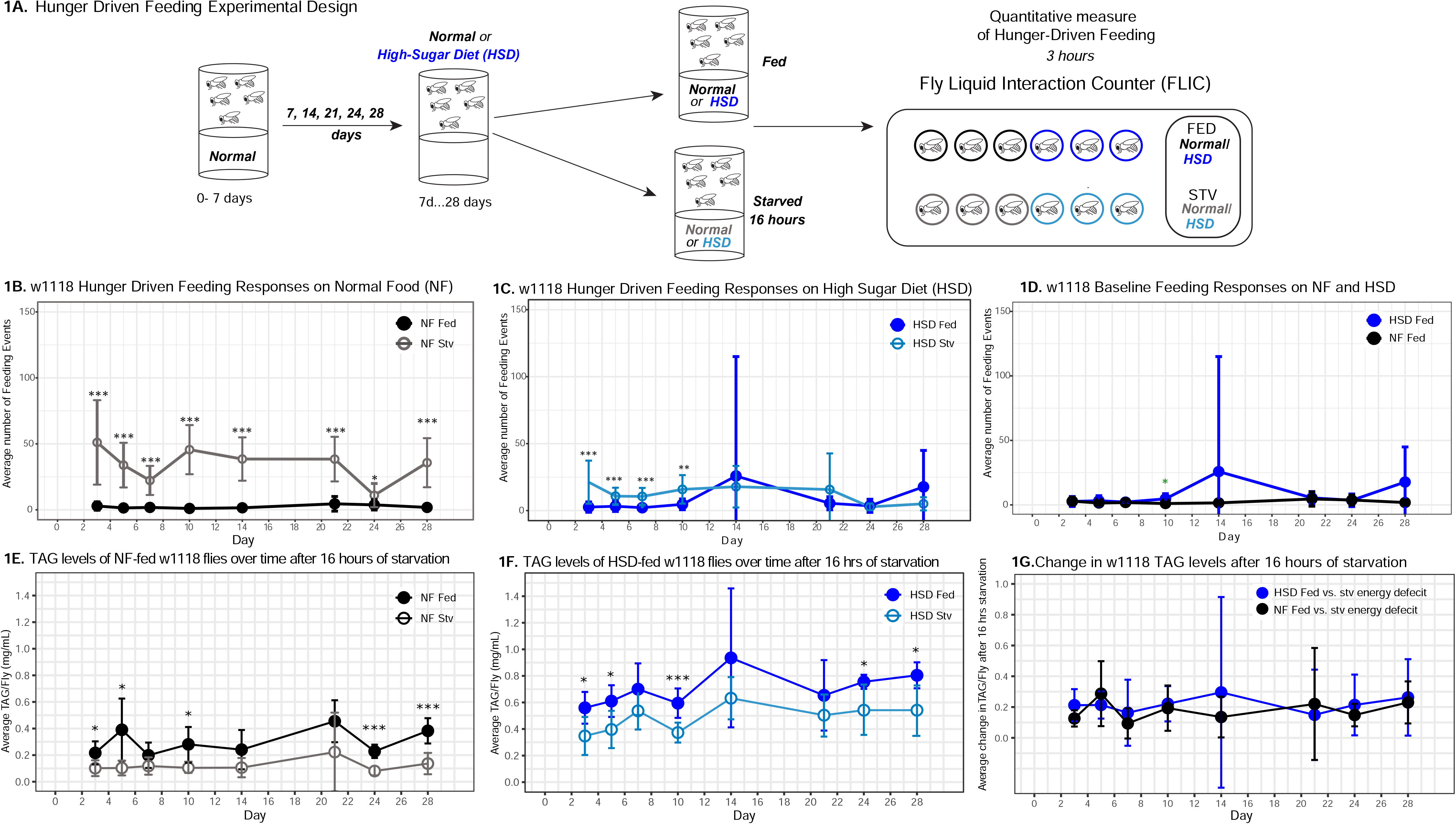
HSD causes progressive loss in hunger-driven Feeding. A) Hunger driven feeding behavior in flies was tested using the schematic in A). After aging flies for 7 days on normal lab food, flies we subject to normal diet or high-sugar diet (30% more sugar in food) for a duration of 3-28 days. For every timepoint, hunger was induced by subjecting flies to starvation (agarose, 0% sucrose media) overnight for 16 hours. Quantitative feeding behavior was monitored in flies subject to NF or HSD using the Fly liquid interaction counter during a 3 hour window immediately after starvation period. Feeding events were recorded and graphed for NF (B, Black), HSD (C, Blue) flies, and the baseline fed states for NF and HSD (D). Average feeding events are plotted for each time point for both *ad libitum* fed (filled dots) and 16 hour starved flies (no fill) with error bars indicating standard deviation (n=18/diet/fed state/day). B) Average feeding events over time for NF flies that were either fed or starved for 16 hours prior to measurement (stv). Note that HDF is maintained throughout the experiment. C) Average feeding events over time for HSD-fed flies that were either fed or starved for 16 hours prior to measurement (stv). Note loss of HDF at day 14. D) Comparison of basal feeding in NF and HSD fed flies over time. (E-F) Average TAG levels per fly of NF (E) and HSD-fed (F) flies at baseline and after 16 hours of starvation. (G) Average change in TAG levels after starvation in NF and HSD flies (n=8-9 for each treatment and timepoint). Two-Way ANOVA with Sidak post-test correction. Asterisks indicate significant changes with *p value <0.05, **p value <0.005, and ***p value <0.0005. Error bars= standard deviation.

We next asked whether flies on HSD are capable of sensing and responding to an energy deficit. Triacylglycerides (TAGs) make up the largest energy reserve and are mobilized when flies are starved^51^. Thus, changes in TAG levels following starvation can be used as a readout for cellular energy-sensing in *Drosophila*^52,53^. Despite having higher baseline levels of TAG (Figure S2), the HSD-fed flies displayed similar levels of starvation-induced TAG breakdown to NF-fed flies throughout the 4-week window (Figure 1E-G). Notably, despite the consistent starvation-induced TAG mobilization, the HDF response only dampens after day 14 of HSD (Figure 1C). Taken together, these results suggest that while chronic HSD treatment leads to increased TAG stores, flies sense starvation and respond to the energy-deficit by mobilizing TAG. Nonetheless, they display a progressive loss of HDF (Figure 1C), suggesting that prolonged HSD exposure results in the uncoupling of feeding behavior from energy-sensing.

### Insulin Resistance and Lipid morphology changes at point of HDF loss

The loss of HDF behavior after 14 days of HSD suggests that this is a critical time point of metabolic disruption. Given that insulin signaling is a major regulator of systemic metabolism^54,55^, and insulin resistance is a hallmark of obesity^56^, we asked whether HSD fed flies display dysfunctional insulin signaling. To answer this, we first measured the amount of *Drosophila* insulin-like peptide 5 (Dilp5) accumulation in the Insulin Producing Cells (IPCs) in the brain. It has been shown that Dilp5 accumulation in the IPCs directly correlates with nutritional status^57^; when flies experience a nutrient-rich environment, Dilp5 levels in the IPCs drop due to increased insulin secretion^58^. Consistent, with previous reports^58^, we found that 14 days of HSD feeding *ad libitum* resulted in decreased Dilp5 accumulation in the IPCs (Figure 2A). To determine whether this was due to a downregulation of Dilps at the transcript level, we analyzed the expression of the IPC-secreted Dilps 2, and 5^59^. We found that 14 days of HSD treatment did not alter the expression of Dilps 2 and 5 (Figure S3A). Taken together, these results (Figure 2A, S3A) suggests that HSD increases Dilp5 secretion.

**Figure 2.**
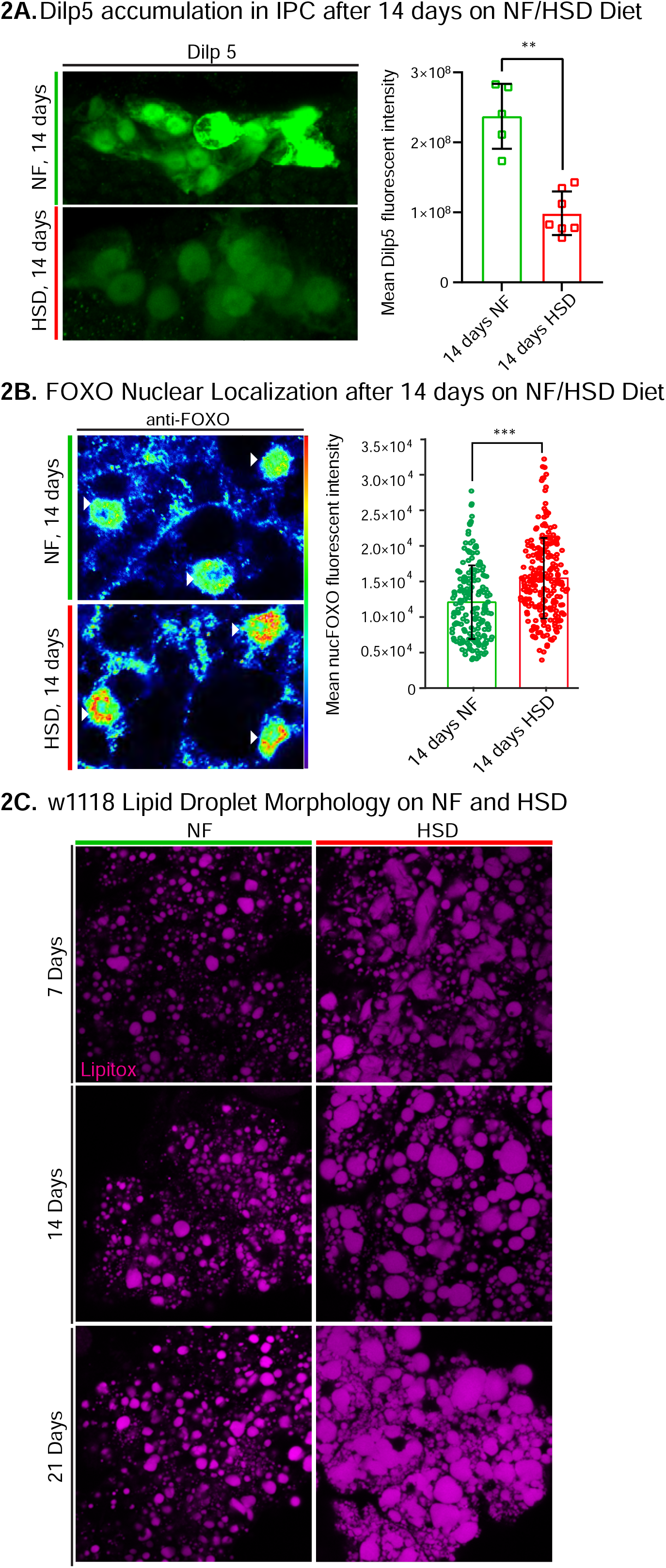
DILP5/FOXO accumulation and Lipid droplet morphology altered under HSD exposure. A, left) Representative confocal images of Dilp5 accumulation in IPCs of NF flies (top panel) and HSD -fed flies at day 14 (bottom panel). A, Right) Mean Dilp5 fluorescent intensity from z-stack summation projections of IPCs from NF and HSD-fed flies. Each square represents a single fly. Unpaired t test with Welch’s correction. B) Representative confocal images of nuclear FOXO accumulation in the fat bodies of NF flies (top panel) and HSD -fed flies at day 14 (bottom panel). The nuclei are marked with anti-lamin (magenta). Arrowheads point to nuclei. B, Right) Mean nuclear FOXO fluorescent intensity from z-stack summation projections of fat bodies from NF and HSD-fed flies. N=each circle represents a nucleus. Two-sided Wilcoxon rank-sum test. Error bars=standard deviation. Asterisks indicate significant changes with *p value <0.05, **p value <0.005, and ***p value <0.0005. C) Representative confocal images of lipid droplets (magenta) across time in the fat bodies of NF and HSD-fed flies.

We reasoned that, as has been previously reported^32^, the changes in insulin signaling may lead to insulin resistance in peripheral tissue. To address this, we measured the forkhead box O (FOXO) nuclear localization in the fat body (Figure 2B). Insulin signaling is activated by the binding of insulin to its cell surface receptor (IR)^60,61^. When activated, IR auto-phosphorylates, which triggers the phosphorylation of multiple downstream targets including the transcription factor FOXO^62,63^. Phosphorylation of FOXO prevents it from entering the nucleus and initiating the gluconeogenic pathway, a starvation response pathway. Thus, FOXO nuclear localization has been used as a proxy to monitor insulin sensitivity^64,65^. Given that the fat body is a major target for insulin signaling^66^, we wondered whether increased insulin signaling in HSD led to insulin resistance. Indeed, we found that 14 days of HSD treatment resulted in elevated nuclear FOXO levels compared to the NF treatment. Notably, this effect was not seen after acute six-hour exposure to HSD (Figure S3B).

Accumulation and enlargement of lipid droplets, the cell’s lipid storage organelles, are associated with insulin resistance^67^. Flies on HSD showed progressively larger, misshapen, and denser lipid droplets compared to flies on NF (Figure 2C). Together, the reduced Dilp5 in the IPCs, increased nuclear localization of FOXO in the fat body, and altered lipid droplet morphology following chronic 14-day HSD treatment suggest insulin resistance.

### Lipidomics on HSD uncovers alterations in whole body phospholipid levels at 14-day HSD

Next, we sought to characterize the progressive changes in the lipidome of flies subjected to HSD using a targeted quantitative lipidyzer (See Methods). Although a short 7-day exposure to HSD did not greatly change the overall concentrations of lipid classes (Figures 3 and S4A), a prolonged 14-day exposure, caused a significant increase in the concentrations of triacyl glycerides (TAGs) and diacyl glycerides (DAGs) (Figure S4A), while free fatty acids (FFAs) were surprisingly lower in HSD fed flies compared to NF flies (See Discussion). Intriguingly, prolonged 14-day exposure to HSD caused a significant increase in the PE and PC levels (Figure 3). We also observed that two other phospholipid classes, lysophosophatidylcholine (LPC) and lysophosphatidylethanolamine (LPE), are downregulated on 14-day HSD, with changes in LPE being statistically significant (Figure 3). We speculate that LPE reduction is a likely effect of upregulated PE synthesis, as LPE serves as a precursor for PE^68^ (See Discussion). Together, these findings indicate that altered phospholipid levels and properties correlate with the point of HDF loss and insulin resistance.

**Figure 3.**
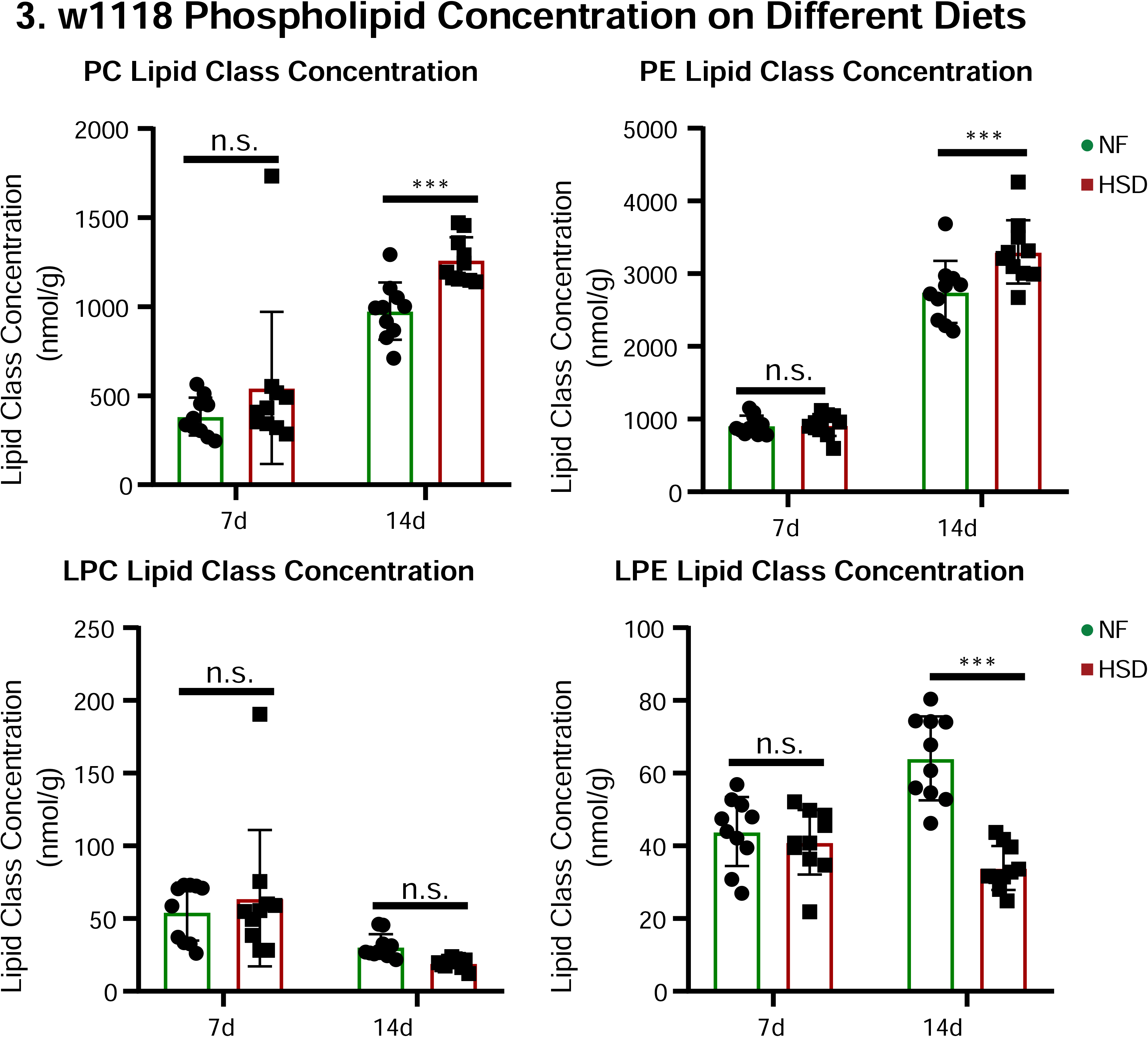
Phospholipids are elevated in whole fly during extended HSD intake. Average concentrations of PC, PE, LPC, and LPE in flies fed subjected to NF (green) or a HSD (red) for 14 days. Lipidomics was performed using a targeted quantitative lipidyzer (Sciex 5500 Lipidyzer). 10 independent biological replicates were used for each diet and each day, with n=10 flies composing 1 biological replicate. Two-way ANOVA and Sidok post-test correction. Asterisks indicate significant changes with *p value <0.05, **p value <0.005, and ***p value <0.0005. Error bars= standard deviation.

### Fat body Pect levels regulate apolipoprotein delivery to the brain

PC and PE are primarily synthesized by the fat body and trafficked in lipophorin particles (Lpps) chaperoned by ApoLpp to other organs, including the brain^69^. ApoLpp is the functional ortholog of human ApoB^69^, and PE-rich ApoII particles have been shown to traffic from fat to brain^69^. Since PE levels are increases on 14 day HSD (Figure 3), we reasoned that we would observe increased ApoII staining in the brain. Using an antibody against ApoII (Figure S5), the fragment of Apolpp that harbors the lipid binding domain, we asked whether 14 days of HSD treatment would increase ApoII delivery to the brain. Surprisingly, we found that though HSD treatment caused an increase in the overall PC and PE lipid levels (Figure 3), the amount of ApoII that chaperones PE to the brain was significantly reduced in an area of the brain proximal to the IPCs (Figure 4A). This observation leads us to propose that fat to brain trafficking of PE via ApoII particles was disrupted by 14-day HSD (See Discussion).

**Figure 4.**
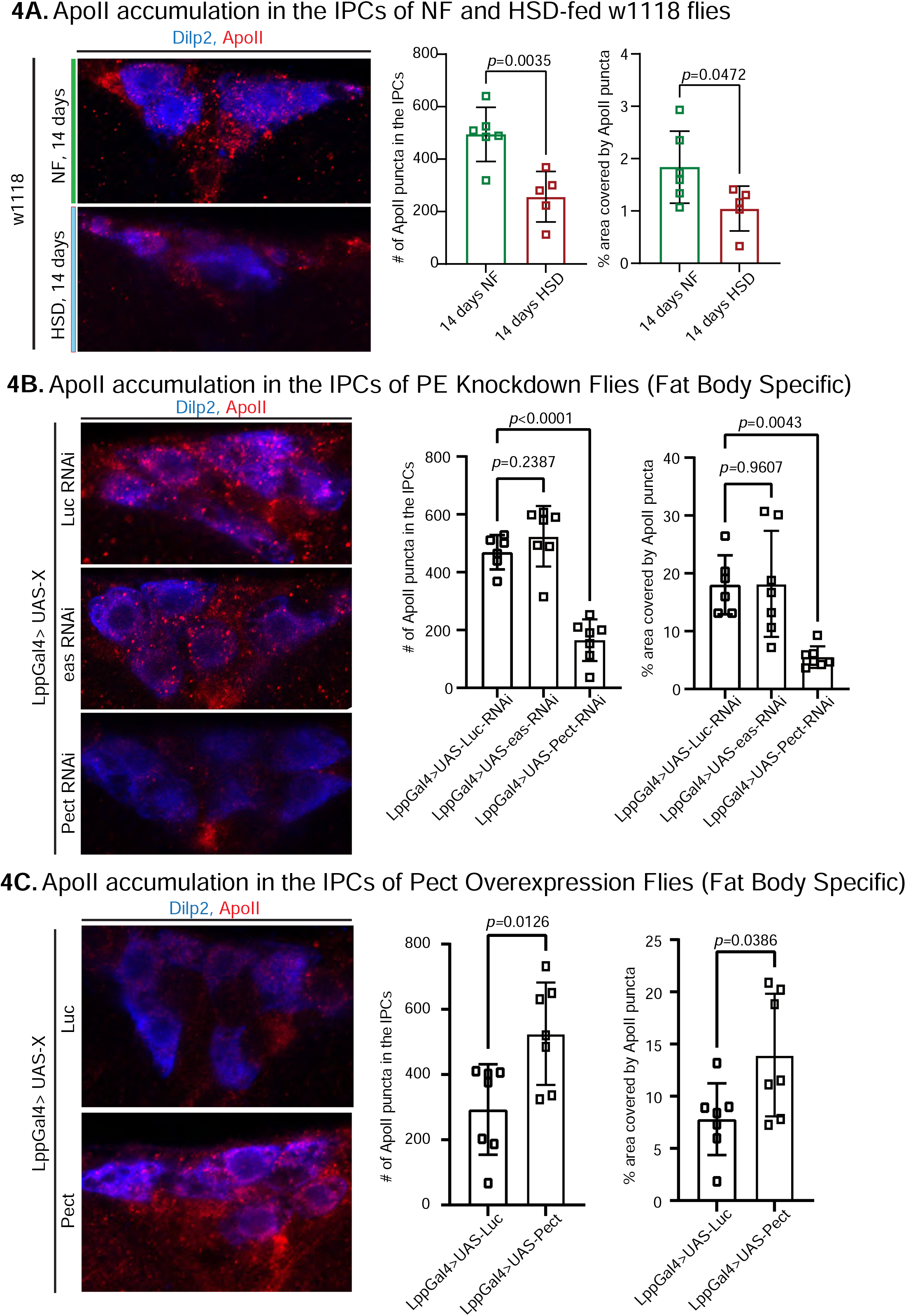
HSD and Knockdown of Pect lead to decreased ApoII levels in the brain. A-C, left) Representative confocal images of ApoII (red) levels in the IPCs (blue) of A) NF and HSD-fed flies at day 14, B) flies with a fat-specific knockdown of Luc, eas, and Pect, and C) flies with a fat-specific overexpression of Luc and Pect. (A-C, right) Mean number of ApoII puncta and % area covered by ApoII puncta in the IPCs of A) NF and HSD-fed flies at day 14, B) flies with a fat-specific knockdown of Luc, eas, and Pect, and C) flies with fat specific overexpression of Luc and Pect. N= each square represents a single fly. Unpaired t-test with Welch’s correction. Asterisks indicate significant changes with *p value <0.05, **p value <0.005, and ***p value <0.0005. Error bars=standard deviation.

ApoLpp is synthesized exclusively by the fat body and that Lpps are enriched in PE^69^. Therefore, we asked whether manipulating PE levels in the fat body via RNAi-mediated knockdown of two important enzymes in the PE biosynthesis pathway, Pect and easily shocked (eas)^24,25,70^, would affect ApoII levels in the brain (see qPCR validation in figure S6A). We found that while eas did not alter the levels of ApoII in the brain, Pect, the rate-limiting enzyme of PE biosynthesis, caused a reduction in ApoII levels like that of HSD (Figure 4B). Conversely, fat body specific Pect overexpression increased ApoII levels in the brain (Figure 4C). Given that HSD and Pect knockdown caused a similar effect on ApoII levels in the brain, we predicted that Pect mRNA levels would be low in the HSD-fed flies. To our surprise, we found that 14 days of HSD caused a 200-270-fold rise in Pect mRNA levels that fell sharply by day 21 (Figure S6B) compared to the modest ∼7-fold increase in the Pect OE flies (Figure S6A, right). This suggests that extreme deregulation of Pect levels, either up or down, may affect PE levels and consequently their carrier ApoII (See Discussion). This interpretation is consistent with previous studies linking Pect dysregulation to abnormal lipid metabolism and signaling^26–28^.

### Fat body specific Pect knockdown shows FOXO accumulation and lipid morphology changes similar to HSD

Given that PE homeostasis is disrupted under HSD (Figure 3), and that coincides with signs of insulin resistance in the fat body (Figure 2B-C), we asked whether fat body-specific knockdown of Pect affects insulin sensitivity. To answer this, we measured the nuclear FOXO levels of fat body specific Pect knockdown flies (Figure 5A). We found that like the HSD-treated flies (Figure 2B), Pect knockdown flies exhibit increased nuclear localization of FOXO. Given that lipid droplets serve as a repository for lipids^71^, the building blocks of phospholipids, we reasoned that knocking down Pect will lead to excessive accumulation of lipids and result in enlarged lipid droplets resembling flies on HSD. Indeed, a qualitative view of lipid droplets in Pect knockdown flies showed larger and more clustered lipid droplets than in control (Figure 5B). These data suggest that Pect is important for mediating insulin sensitivity in the fat body, and its loss leads to DIO-like phenotypes.

**Figure 5.**
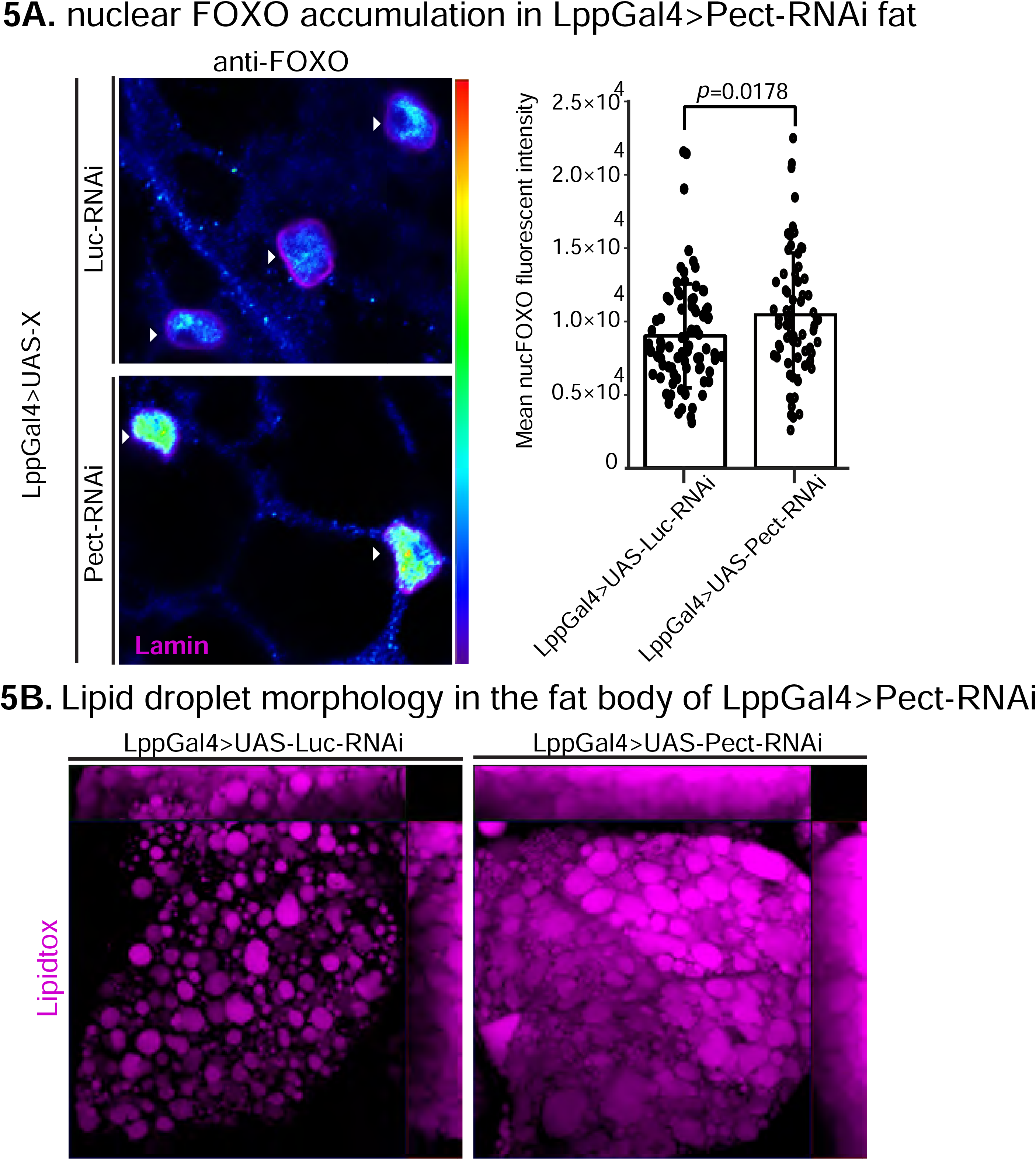
Pect knockdown in fly fat alters FOXO nuclear accumulation and lipid droplet morphology. A, left) Representative confocal images of Lpp-Gal4>UAS-Luc-RNAi and Lpp-Gal4>UAS-Pect-RNAi fly fat stained for anti-FOXO (blue to green, intensity-based) and lamin (pink) on day 14. Arrowhead points to the nucleus. (A, left) Mean nuclear FOXO fluorescent intensity of fat bodies from control (LppGal4>UAS-Luc-RNAi) and fat-specific Pect knockdown flies (LppGal4>UAS-Pect-RNAi). N=each circle represents a nucleus. Two-sided Wilcoxon rank-sum test. Error bars=standard deviation. Asterisks indicate significant changes with *p value <0.05, **p value <0.005, and ***p value <0.0005. Error bars= standard deviation. B) Representative confocal images of lipid droplets (magenta) in the fat bodies of control (LppGal4>UAS-Luc-RNAi) and fat-specific Pect knockdown flies (LppGal4>UAS-Pect-RNAi). Note that lipid droplet morphology in the fat-specific Pect knockdown flies resembles that of wild-type flies on HSD.

### Pect Knockdown in the Fat Body alters Whole Body Phospholipid Concentrations

Next, we sought to determine the impact of Pect on the lipidome of adult flies. To do this, we compared the lipidomic profile of whole flies expressing Pect-RNAi using a fat specific driver (*Lpp-Gal4*) against a control-RNAi (luciferase-RNAi; Figure 6). We found that knocking down Pect in the fat body slightly reduced whole body PE levels (Figure 6A). This is consistent with a previous study showing that *pect* null mutants did not display significant alterations in the levels of PC and PE^25^. However, consistent with the same study^25^, we observed that specific PE species showed significant alterations, with PE 36.2 displaying a significant downregulation (Figure 6B) (See Discussion). Furthermore, while other major lipid classes did not significantly deviate from control (Figure S8), we noted a striking and significant increase in two minor classes of phospholipids LPC and LPE (Figure 6A). Given that LPE can serve as a precursor for PE via the exogenous lysolipid metabolism (ELM) pathway^68^, we speculate that in the absence of the rate-limiting enzyme for PE synthesis, there is an elevation in LPE levels (See Discussion). We also performed the same analysis in a Pect overexpression fly line, but found no change in any of the lipid classes compared to control (Figure S9). Together, this suggests that fat-specific knockdown of Pect is sufficient to cause a reduction in certain PE species and upregulation of the minor phospholipid classes, especially the PE precursor class of minor phospholipid LPE. Hence, we conclude that fat-specific Pect disruption results in phospholipid composition imbalance.

**Figure 6.**
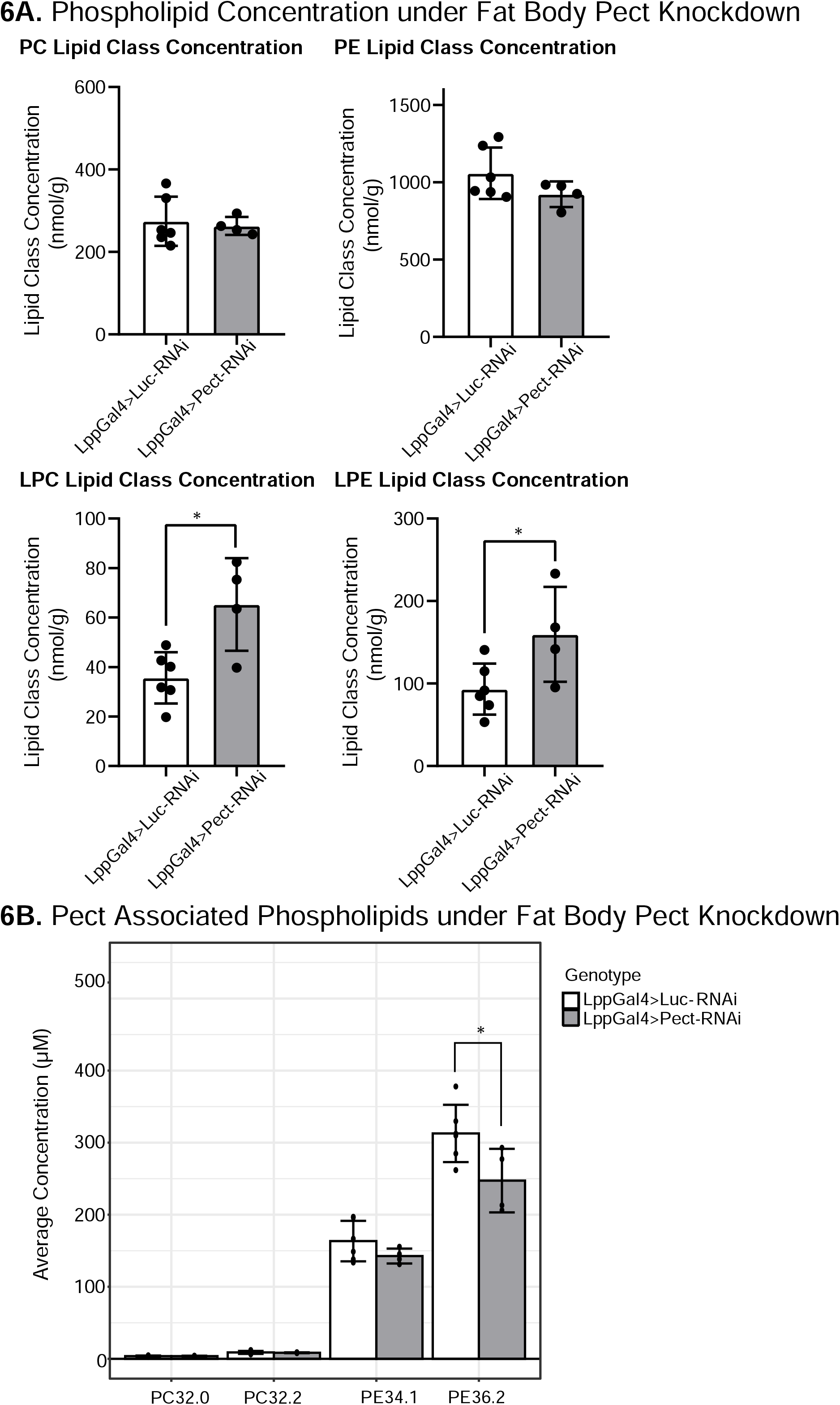
Manipulating Pect levels in the fat body alters phospholipid profile. A) Average concentrations of PC, PE, LPC and LPE lipid classes in LppGal4>UAS-Pect-RNAi flies compared to control under 7 day NF conditions. B) Depicts the concentration of Pect associated phospoholipids in LppGal4>UAS-Pect-RNAi flies compared to control. Lipidomics was performed using a targeted quantitative lipidyzer (Sciex 5500 Lipidyzer). 4-6 independent biological replicates were used for each genotype, with n=10 flies composing 1 biological replicate. Unpaired t-test with Welch’s correction. Asterisks indicate significant changes with *p value <0.05, **p value <0.005, and ***p value <0.0005. Error bars= standard deviation, points= individual replicate values.

### Pect knockdown in the fat modulates hunger driven feeding behavior

We next sought to determine whether phospholipid imbalance would impact feeding behavior. Given that the fat body is a major source of phospholipids synthesis^69^, we asked whether disrupting PE and PC synthesis would impact HDF behavior in NF and HSD feeding conditions. To answer this, we knocked down key enzymes in the PE and PC biosynthesis pathways in the fat body via RNAi and assessed HDF behavior in response to starvation when flies are fed NF (Figure 7A). We found that knocking down the PC-biosynthesis enzyme Pyct2 did not affect HDF (Figure 7A). Similarly, knock down of eas, which initiates the early enzymatic reaction of PE biosynthesis^70^ did not have an effect of HDF (figure 7A). However, knocking down the rate-limiting enzymes in the ER (Pect) and mitochondria (PISD)-mediated PE biosynthesis pathways^72^ led to a diminished HDF response (Figure 7A). Interestingly, knock down of eas, PISD, and Pect all led to loss of HDF after 14 days of HSD treatment (Figure 7A). This suggests that, unlike PC, PE plays a critical role in regulating HDF behavior in both baseline and HSD conditions. Indeed, fat body-specific Pect overexpression (Pect OE) was sufficient to rescue HDF loss in HSD-fed flies (Figure 7B). Pect OE flies maintain their starvation response after 14 day of HSD exposure, a critical time point for HDF breakdown as seen in the wild type flies (Figure 1C and 7B). Together, these data indicate that Pect activity and the maintenance of proper phospholipid homeostasis in the fat body is critical for maintaining an HDF response.

**Figure 7.**
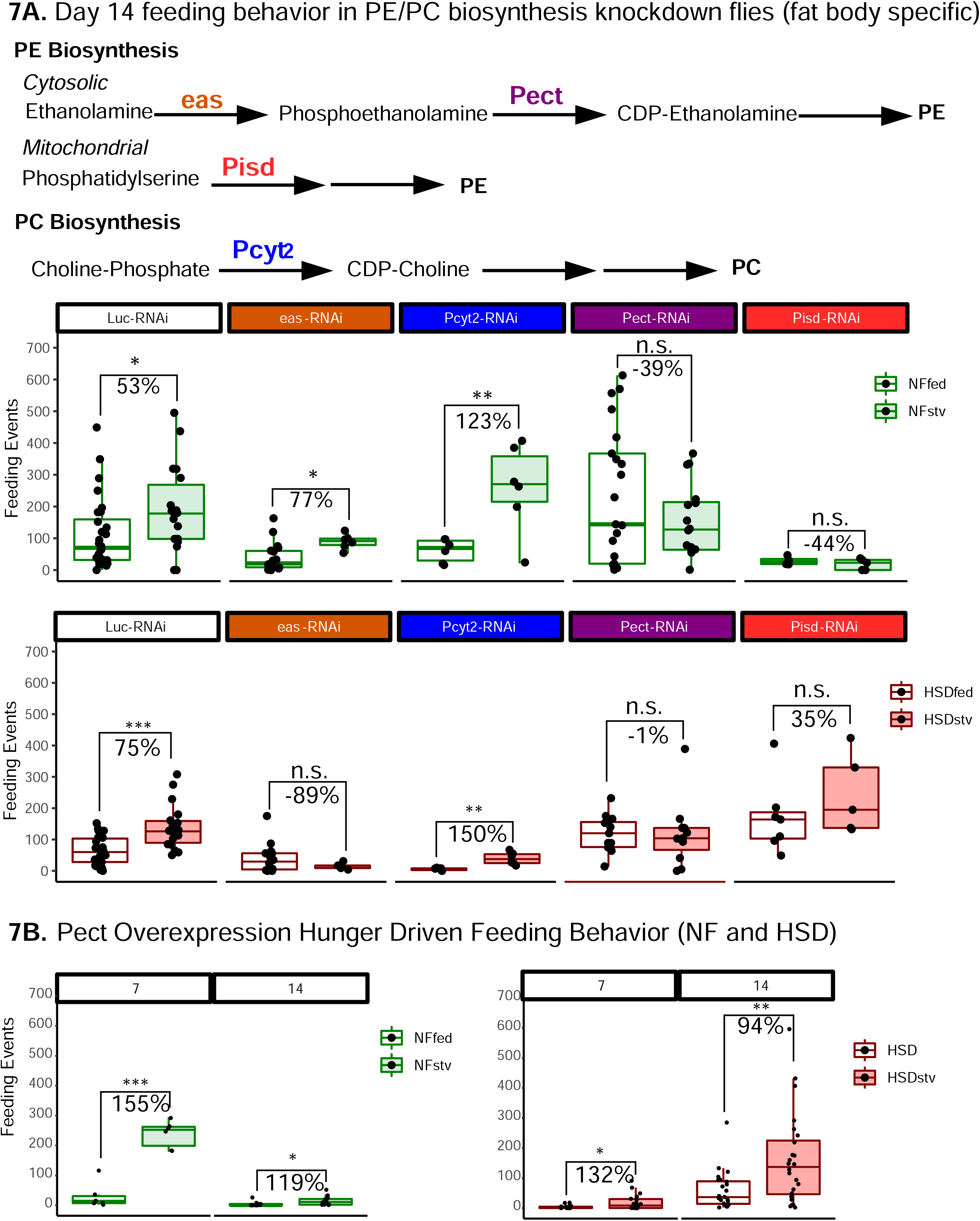
Manipulating Pect levels in the fat body alters hunger Driven Feeding behavior. A) A simplified schematic of the PE and PC biosynthesis pathway. eas and Pect are enzymes in the PE biosynthesis pathway, whereas PCYT2 is an enzyme in the PC biosynthesis pathway. B, top) Average fly feeding events over time for NF flies that were either fed or starved for 16 hours prior to measurement (stv). Note that HDF is maintained throughout the experiment. A, bottom) Average feeding events over time for HSD-fed flies that were either fed or starved for 16 hours prior to measurement (stv). Note loss of HDF at day 14. Each dot denotes an individual fly. Two-Way ANOVA with Sidak post-test correction. Asterisks indicate significant changes with *p value <0.05, **p value <0.005, and ***p value <0.0005.. Error bars= standard deviation. B) Average feeding of fed and starved Lpp-Gal4>UAS-eas-RNAi (orange), Lpp-Gal4>UAS-PCYT2-RNAi (blue), Lpp-Gal4>Pect-RNAi (purple), and Lpp-Gal4>PISD-RNAi (red)after 14 days of NF (B,top) or HSD (B, bottom).

## Discussion

In recent decades, the prevalence of obesity has reached pandemic proportions^1^. Increased consumption of excess sugars is considered a major contributing factor^73^. HSD-induced obesity has been shown to lead to a plethora of physiological dysfunctions including insulin resistance and type 2 diabetes^74^. Several studies have shown a link between chronic sugar consumption and altered hunger perception^75,76^. Although the neuronal circuits governing hunger and hunger-driven feeding behavior have been well studied^4^, less is known about the impact of adipose tissue dysfunction on feeding behavior. Using a *Drosophila* model of DIO, we show that phospholipids, specifically PE, play a crucial role in maintaining hunger-driven feeding behavior.

The *Drosophila* model organism has been shown to be a relevant model for mimicking the conditions of human diet-induced obesity and insulin resistance^34^. Studies from Dus and Ja et al. have previously performed measurements on taste preference, feeding behavior/intake, survival, etc. using a high sugar diet-induced obesity model and have found much in common with their mammalian counterparts^36,42^. However, the longest measurement of adult feeding behavior has been capped at 7 days^47^. A recent study by Musselman and colleagues has performed analysis of lipidome on 3-week and 5-week HSD in a tissue-specific manner^77^ and identify changes in neutral fat stores. Their important study provide a snapshot of what happens to lipidome changes at late-stage HSD, and added further credence to the idea that neutral fat stores – TAG and DAG – are upregulated on HSD in the cardiac tissue ^77^. In our study by capturing systemic lipid changes on DIO at a time-point (14-day HSD) where adult flies fed HSD begin to lose homeostasis, we have sought to provide insights into how HSD exposure uncouples energy-sensing from feeding behavior. By doing so, we have uncovered a critical requirement for the activity of the rate-limiting PE enzyme Pect within the fat tissue in controlling hunger-driven feeding. Furthermore, based on our findings, we propose that maintaining a tight PE and LPE balance is an important factor in regulating hunger-driven feeding response.

### HSD leads to insulin resistance and a progressive loss of hunger-driven feeding behavior

Changes in feeding behavior in both vertebrates and invertebrates occur via communication between peripheral organs responsible for digestion/energy storage and the brain^78^. This communication is facilitated by factors that provide information on nutritional state^78^. One example of such a factor is leptin, which is released from the adipose tissue and acts on neuronal circuits in the brain to promote satiety^79,80^. While Leptin has long been studied as a satiety hormone, recent work in mice and in flies suggests that a key function of leptin and its fly homolog upd2 is to regulate starvation response^10,79,81,82^. Indeed, we have previously shown that exposing flies to HSD alters synaptic contacts between Leptin/Upd2 sensing neurons and Insulin neurons, however, it resets within 5-days^10^. This suggests that homeostasis on surplus HSD diets beyond 5 days is maintained by yet-to-be-defined mechanisms.

To delineate how HSD alters the starvation response, we analyzed feeding behavior over time. We found that under normal diet conditions, flies display a clear response to starvation in the form of elevated feeding that we termed “hunger-driven feeding (HDF)” that was independent of age (Figure 1B). In contrast, chronic exposure to HSD led to a progressive loss of HDF that began on day 14 (Figure 1C). It could be argued that loss of HDF is simply due to an elevation of TAG storage in HSD fed flies, thus losing the need to feed on starvation. TAG levels were found to be significantly elevated at all timepoints compared to NF (excepting day 21, Figure S2), thus we would expect all flies under HSD diet to show no HDF response by day 5 onward. However, our data does not lend support the argument as we find that HDF is sustained in HSD-fed flies until day 14, despite baseline levels of TAG being significantly elevated from 5 days of HSD treatment onward. Moreover, the reduced HDF response is not a result of increased *ad libitum* baseline feeding on HSD, as HSD-fed flies consume food at levels similar to NF-fed flies (Figure 1D). Hence, we favor the conclusion that despite elevated TAG levels (Figure S2), HSD-fed flies are motivated to seek food on starvation up until Day 14 (Figure 1C).

It is to be noted that the HDF response of HSD-fed flies (Figure 1C, Days 3-10) is of lower order of magnitude than the NF-fed flies. This suggests that that in addition to sensing an energy deficit and mobilizing fat stores (Figure 1F, 1G, S1), HSD fed flies calibrate their starvation-induced feeding to compensate only for the lost amount of fat. Overall, this suggests that flies have a remarkably fine-tuned ability to coordinate food-intake with nutrient store levels. Nonetheless, beyond Day 14, this capacity of flies to couple energy sensing and food intake is lost, as evinced by the loss of HDF beyond Day 14 HSD (Figure 1C, S1). Strikingly, despite a prolonged 4-week exposure to HSD, HSD-fed flies can sense an energy deficit at 4-weeks given that they can effectively mobilize fat stores in response (Figure 1F, 1G). Taken together, our experiments suggest that prolonged exposure to HSD leads to an uncoupling of energy deficit and feeding behavior.

Notably, there are striking differences and similarities between fly and mammalian DIO models). While mice show linear weight gain on obesogenic diets, the flies’ rigid exoskeleton limits their capacity to store TAG beyond a certain point ^83–85^ (Figure S2). However, similar to mammals, we find that prolonged exposure to HSD, which is strongly associated with phospholipid dysregulation^16–18^, leads to reduced insulin sensitivity^17,32^. We show that the levels of Dilp5, the fly’s insulin ortholog, are reduced in the IPCs of HSD-fed flies (Figure 2A), however we do not detect a decrease in Dilp5 or Dilp2 mRNA levels (Figure S3A), indicative of increased insulin secretion on HSD, and similar to what has been previously reported by the Leopold lab for HSD fed larvae^58^. Consistent with the idea that 14-day HSD triggers insulin resistance, we observe elevated FOXO nuclear localization in the fat bodies of the HSD-fed flies (Figure 2B), despite a likely increase in Dilp5 secretion on HSD (Figure 2A). Again, these findings align with mammalian studies showing that dysregulated FOXO signaling is implicated in insulin resistance, type 2 diabetes, and obesity^65^.

### HSD and fat-specific PECT-KD causes changes to phospholipid profile

Changes in the lipidome are strongly correlated with insulin resistance and obesity^86^. However, less is known about how the lipidome affects feeding behavior. To this end, we analyzed the lipid profiles of NF and HSD-fed flies over time (Figure 3). As expected, exposure to HSD increased the overall content of neutral lipids compared to the NF flies with TAGs and DAGs increasing the most, which is consistent with other DIO models^32,36,47^. Surprisingly, we noted that 14 days of HSD treatment caused a decrease in FFAs and a rise in TAGs and DAGs (Figure S4A). We speculate that this reduction in FFA maybe due to their involvement in TAG biogenesis^87^. We were interested to see if the decrease in FFA correlated to a particular lipid species, as PE and PC are made from DAGs with specific fatty acid chains. However, further analysis of FFAs at the species level did not reveal any distinct patterns. The majority of FFA chains decreased in HSD, including 12.0, 16.0, 16.1, 18.0, 18.1, and 18.2 (Figure S4B). This data was more suggestive of a global decrease in FFA, likely being converted to TAG and DAG, rather than a specific fatty acid chain being depleted.

On 14-day HSD, when HDF response begins to degrade (Figure 1C, S1), Phosphatidylethanolamine (PE) and Phosphatidylcholine (PC) levels rise dramatically, whereas lysophosphatidylethanolamine (LPE) significantly decreases. Interestingly, similar patterns of phospholipid changes have been associated with diabetes, obesity, and insulin resistance in clinical studies^16–18,24^, yet no causative relationship has been established^16–18^. On future examination, we find that PC balance appears dispensable for maintenance of HSD-response, but both the mitochondrial and cytosolic PE pathway seem to be critical for HDF response (Figure 7). PE is synthesized by multiple pathways. Studies have shown that in addition to the mitochondrial PISD^88^ and cytosolic CDP-ethanolamine Kennedy pathway^89,90^, PE can be synthesized from LPE^68^. This pathway is named the exogenous lysolipid metabolism (ELM) pathway, can substitute for the loss of the PISD pathway in yeast and requires the activity of the enzyme lyso-PE acyltransferase (LPEAT) that converts LPE to PE^91^. In this study, we noted that PE levels were upregulated on HSD, while LPE levels were downregulated (Figure 3). In contrast, fat-specific Pect-KD caused PE levels to trend downward, whereas LPE was upregulated (Figure 6A). Though the level changes for PE and LPE *per se* are contrasting between 14-day HSD lipidome and Pect-KD, there is a critical significant commonality in that under both states there is an imbalance of phospholipids classes PE and LPE. Hence, we propose that maintaining the compositional balance of phospholipid classes PE and LPE is critical to hunger-driven feeding and insulin sensitivity.

It is worth noting that the role of the minor phospholipid class LPE remains obscure. In our study, we observe that the LPE imbalance occurs on both prolonged HSD exposure and when fat body Pect activity is disrupted. This suggests that LPE balance is likely to play a role in insulin sensitivity and the regulation of feeding behavior. We anticipate that this observation will stimulate interest in studying this poorly understood minor phospholipid class. In future work, it would be interesting to test how the genetic interactions between the enzyme that converts LPE to PE, called LPEAT^91^ and Pect manifest in organismal hunger-driven feeding. Specifically, it will be interesting to ask whether reducing or increasing LPEAT will restore PE-LPE balance to improve the HDF response in HSD-fed flies and Pect-KD. In sum, future studies should explore how LPE-PE balance can be manipulated to affect feeding behaviors.

In addition to changes in phospholipid classes (Figure S7C), we found that HSD caused an increase in the concentration of PE and PC species with double bonds (Figure S4C and S4D). Double bonds create kinks in the lipid bilayer, leading to increased lipid membrane fluidity which impacts vesicle budding, endocytosis, and molecular transport^14,92^. Hence it is possible that a mechanism by which HSD induces changes to signaling is by altering the membrane biophysical properties, such as by increased fluidity, which would have a significant impact on numerous biological processes including synaptic firing and inter-organ vesicle transport. Consistent with this idea, despite the presence of higher amounts of PE lipid species on HSD (Figure 3), we observe significant reduction in the trafficking of Apo-II positive lipophorin particles from fat tissue to the brain (Figure 4A). Hence in the future, it would be important to explore how HSD alters the membrane environment of adipose tissue and the brain to impact behavior.

### The implications of the relationship between Pect levels and HSD

The lipidome analysis revealed that many of the lipid species and classes that increase under HSD are regulated by Pect, the rate-limiting enzyme of the PE biosynthesis pathway. Because PCYT2, the human ortholog of Pect, has been associated with BMI in GWAS studies^16^, we reasoned that it may play a role in regulating insulin sensitivity and feeding behavior. Supporting a critical role for the activity of Pect in the fat body for maintenance of systemic homeostasis, we find that Pect disruption in the fat body reduces the fat-brain transfer of ApoII-positive lipophorins (Figure 4B), reduces insulin sensitivity (Figure 5A), and displays abnormal lipid droplet morphology that qualitatively phenocopies the prolonged 14-day HSD (Compare Figure 5A to 2C). Significantly, Pect-KD phenocopies the loss of hunger-driven feeding response observed in 14-day HSD. Given all this, we expected that Pect levels would be depressed on 14-day HSD, but we were surprised to find that 14-day HSD causes a 200-250 upregulation of Pect mRNA. To resolve this, we performed lipidomic analyses on Pect-KD to understand how that might compare to 14-day HSD. We noted that on whole body lipidomics performed on flies with Pect-KD, PE was reduced though not significantly (Figure 6A), PC was unchanged, but the minor phospholipid classes LPE and LPC were significantly upregulated. This change in Pect-KD phospholipidome contrasts in 14-day HSD in that LPE is significantly reduced and PE is significantly upregulated (Figure 3). These differences are consistent with our observation that Pect is upregulated on HSD (figure S6B). Taken together, the Pect-KD lipidome and 14-day HSD lipidome display opposing effects on LPE and PE levels. Nonetheless, the common feature is the dysregulation of Pect levels and phospholipid imbalance. Hence, we interpret this to mean that extreme dysregulation of Pect levels impacts insulin sensitivity and HDF response. Also, we note that while over-expression of Pect cDNA in the fat-body does not alter phospholipid balance (Figure S9), it improves HDF on HSD (Figure 7B). Although this may appear inconsistent, it is critical to note that over-expression of Pect cDNA using UAS/Gal4 only increases Pect mRNA expression by 7-fold (Figure S6A), whereas HSD causes its upregulation by 250-fold (Figure S6B). Hence, we speculate that an increased ‘basal’ level of Pect such as by that provided by a cDNA over-expression in fat, may be protective to the negative effects of HSD (Figure 7B) without affecting overall phospholipid levels (Figure S9), but extreme upregulation Pect on HSD affects the PE and LPE balance (Figure 3).

### Comparison of our fact specific Pect-KD results with prior work

A prior study by Tsai and colleagues^25^ have shown that Pect is required in *Drosophila* photoreceptors for synaptic vesicle pool maintenance^25^. Pect loss of function (LOF) mutations (alleles *LL0632*5 and *omb593*) result in lethality, and hence, in this prior study lipidomic analysis of Pect mutant retinas was carried out. It was identified that when Pect was mutated in fly retinas, the overall proportions of PC and PE species were not significantly changed (81% PC species and 19% PE species); this is consistent with what we observe in global PC and PE levels under a fat-specific Pect KD (Figure 6A). However, loss of Pect activity caused a significant shift in the relative levels of a specific subset of PC and PE species (See Figure 3, Tsai et al.^25^). Of critical relevance to this study, PE 36.2 was downregulated in Pect mutant retinas. We observe the same PE lipid species is significantly downregulated in the whole body lipidome of fat-specific Pect-KD (Figure 6b).Notably, not all pect-dependent PE and PC lipid classes that were identified in the *pect null mutants* were altered in our study, for instance, we did not observe a significant increase in PC 32.2. We attribute these differences to both the context specificity i.e., retina versus whole body lipidomics, as well as the fact that we are assessing whole body lipidome while only impacting Pect activity in fat tissue. It is important to note that we also see Pect associated species disrupted on flies after 14 days of HSD, albeit different species are significantly impacted (PC 32.0 and PC 32.2, Figure S7.). Also, we would like to highlight that since Pect LOF alleles (*pect*^LL06325^ and *pect^omb593^* are lethal, and mosaic analysis is incompatible with studying effects on whole animal physiology, we have relied on fat tissue-specific knockdown of Pect using RNAi.

### Pect activity in fat and its impact on fat-brain communication

To explore the idea that fat-brain communication may be perturbed under HSD and Pect knockdown, we chose to examine a fat-specific signal that is known to travel to the brain. ApoLpp chaperones PE-rich vehicles called lipophorins that traffic lipids from fat to all peripheral tissues including the brain^69^. Consistent with this hypothesis, we observed reduced ApoII levels in HSD and Pect knockdown flies (Figure 4A-B). It has been shown by Eaton and colleagues that lipophorins regulate systemic insulin signaling by impinging on a subset of neurons in the brain^93^. However, it is currently unclear if ApoII is the direct regulator of feeding or whether it is ferrying signaling molecules along with PE/PC lipids. Similar to HSD-fed flies, Pect knockdown flies display dampended hunger-driven feeding even on a normal diet by day 14. This change seemed specific to the PE biosynthesis pathway, as PISD, the critical enzyme for mitochondrial PE biosynthesis in flies, showed a similar impact on HDF. However, we did not observe this effect with Pcyt2, the key enzyme for PC biosynthesis. eas, a non-rate limiting component of PE biosynthesis, showed a decreased effect compared to Pect, only showing changes in HDF in the normal fed. This suggests only a major disruption in PE biosynthesis is required to create the impacts seen in chronic HSD feeding.

## Conclusion

We find that chronic HSD exposure results in significant alterations in the phospholipid profile at the onset of insulin resistance and the loss of hunger-driven feeding behavior. Thus, we have uncovered a role for the phospholipid enzyme Pect as an important component in maintaining hunger-driven feeding. Furthermore, the evidence we present here provides a causative link between phospholipid dysregulation and feeding behaviors. Future work should explore the precise mechanism of how Pect and the associated disruption in phospholipid homeostasis can impact adipose tissue signaling. Nevertheless, this study lays the groundwork for further investigation into Pyct2/Pect as a potential therapeutic target for obesity and its associated co-morbidities.

## Supporting information

Supplemental Table 1- 7-day NF and HSD lipidomics

Supplemental Table 2- 14-day NF and HSD lipidomics

Supplemental Table 3- Fat-specific PECT RNAi and Over-expression lipidomics

## Acknowledgements

We would like to thank Dr. Pierre Leopold for generously donating the FOXO antibody used in this manuscript. We are grateful to Dr. Thomas Clandinin for gifting the UAS-PECT transgenic flie and late Dr. Susan Eaton for the generous gift of the Lpp-Gal4 flies. We would also like to thank the team at the Northwest Metabolomics Research Center for their support in lipidomic profiling. This work was possible due to grants awarded to AR from NIGMS (GM124593), and New Development funds from Fred Hutch Cancer Center. KPK was supported by the NIH Chromosome Metabolism and Cancer Training Grant (T32CA009657) and is by the NSF Post-Doctoral Research Fellowship (NSF Award #2109398). Genomic reagents from the DGRC which is funded by NIH grant 2P40D010949 were used in this study. Stocks obtained from the Bloomington Drosophila Stock Center (NIH P40OD018537) and Transgenic RNAi Resource Project (NIGMS R01 GM084947 and NIGMS P41 GM132087) were used in this study. We are immensely grateful to the three anonymous peer-reviewers for their incisive, constructive, and balanced feedback that significantly contributed to improving the clarity of this study.

## Author Contributions

Conceptualization: AR

Investigation: MEP, CES, AEB, KPK, MAA, ZHG

Formal analysis/Data Curation: AEB, KPK, ZHG, CES, MAA

Visualization: KPK, MAA, JD

Supervision: AR

Writing- original draft: KPK

Writing- review and editing: AR, MAA, KPK

Funding Acquisition: AR

## Methods

### Animals used and Rearing conditions

The following strains were used in this manuscript: w1118, PISD-RNAi (Bloomington #67763), Pect-RNAi (Bloomington #67765), eas-RNAi (Bloomington #38528), Pcyt2-RNAi (Bloomington #67764), Lpp-Gal4 on X (P.Leopold/ S. Eaton.^57^), Luciferase-RNAi (JF01355), and w;;UAS-Pect III (Flybase ID:FBal0347227, generously donated by Clandinin lab), UAS-HA-Apolopp-myc (generously donated by S.Eaton)^94^. Flies were housed in 25°C incubators. In all experiments, only adult male flies were used. Flies were sexed upon eclosion and place on normal diet, a standard diet containing 15g yeast, 8.6g soy flour, 63g corn flour, 5g agar, 5g malt, 74mL corn syrup per liter, for 7 days. Anesthesia using a CO2 bubbler was used for initial sexing, then never used for the remainder of the experiment. After 7 days, flies were either maintained on normal diet or moved to a high sugar diet (HSD), composed of the same composition as normal diet but with an additional 300g of sucrose per liter (30% increase) for the length specified in the figures (typically 7 or 14 days). For measurements of hunger driven feeding, a portion of flies from each diet were placed on starvation media (0% Sucrose/1% agar) for 16 hours prior to the experiment.

### Immunostaining

Immunostaining of adult brains and fat bodies were performed as previously described^10,95^. Tissues were dissected in ice cold PBS. Brains were fixed overnight in 0.8% Paraformaldehyde (PFA) in PBS at 4°C, and fat bodies were fixed in 4% Formaldehyde in PBS at room temperature. Following fixation, tissues were washed 5 times in 0.5% BSA and 0.5% Triton X-100 in PBS (PAT). Tissues were pre-blocked in PAT+ 5% NDS for 2 hours at room temperature, then incubated overnight with the primary antibody in block at 4°C. Following incubation, tissues were washed 5 times in PAT, re-blocked for 30 minutes, then incubated in secondary antibody in block for 4 hours at room temperature. Samples were washed 5 times in PAT, then mounted on slides in Slow fade gold antifade. Primary antibodies were as follows: chicken anti-Dilp2 (1:250); rabbit anti-Dilp5 (1:500); rabbit anti-ApoII (1:500), rabbit anti-FOXO (1:500, gift from Leopold Pierre); mouse-anti-Lamin (1:100; ADL67.10 DSHB). Secondary antibodies from Jackson ImmunoResearch (1:500) include goat anti-rabbit Alexa 647; goat anti-rabbit Alexa 488; donkey anti-chicken Alexa 647; goat anti-rabbit Alexa 594. Lipid droplets were stained with lipidtox (1:500, Thermofischer) overnight at room temperature. Images were captured with Zeiss LSM 800 confocal system and analyzed with ImageJ.

### Image analysis

Quantification of the number of puncta and percentage area occupied by ApoII immunofluorescence was done using ImageJ. Maximum-intensity projections of z-stacks that spanned the entire depth of the IPCs at 0.3 um intervals were generated. A region of interest was manually drawn around the IPCs and a binary mask for the ApoII channel was created using automated Moments thresholding values, which was followed by watershed post-processing to separate particles. The number of particles and the area fraction were measured using the “analyze particles” function.

The fluorescent intensity of Dilp 5 was measured using Z-stack summation projections that included the full depth of the IPCs. A region of interest around the IPCs was manually drawn and the integrated density values were acquired using ImageJ^96^. To measure nuclear FOXO accumulation, a similar number of confocal stacks were acquired for each tissue sample. For FOXO, image analysis was performed in the MATLAB R2020b environment, and the associated scripts are available at github address. FOXO accumulation in *Drosophila* adult fat cells was assessed by measuring the mean voxel GFP intensity in the nucleus that is delimited by the lamin antibody signal. To generate the 3D nuclear masks, Lamin stacks were first maximally projected along the z-axis, and after global thresholding, basic morphological operations and watershed transforms, the locations of the nuclear centroids in the x-y plane were used to scan the z-stacks and reconstruct the nuclear volume. The accuracy of the segmentation was assessed by manual inspection of random cells. Once the nuclear compartment was reconstructed in 3D, it was used as a volumetric mask to extract intensity values of the FOXO reporter signal, and to compute the mean voxel intensity in the nucleus. Two-sided Wilcoxon rank sum tests were performed to assess the statistical significance of pairwise comparisons between experimental conditions.

### Lipidomics

Whole adult male flies were flash frozen in liquid nitrogen either after 7 or 14 days on normal diet or HSD. 10 flies were used per biological sample and 10 biological replicates were used for each diet and timepoint. For Pect manipulation lipidomics, LppGal4>UAS-Luc-RNAi, LppGal4>UAS-Luc, LppGal4>UAS-Pect-RNAi, and LppGal4>UAS-Pect flies were generated and flash frozen after 7 days on NF. Frozen samples were sent to the Northwest Metabolomics Research Center for targeted quantitative lipid profiling using the Sciex 5500 lipidyzer (see Hanson et al. for detailed methods^97^).

### Feeding Behavior

For hunger-driven feeding analysis, age-matched w1118 flies were given a normal diet or HSD for 5, 7, 14, 21, 24, and 28 days after an initial 7 days of development on a normal diet. All other experiments were performed for 7-day or 14-day durations. 16 hours prior to feeding behavior assessment, half of the flies from each treatment were moved to starvation media. Individual flies were placed in a single well of fly liquid-food interaction counter (FLIC) and supplied with a 0% sucrose liquid diet. For feeding behavior involving Gal4>UAS manipulation, a 1% sucrose was substituted for 16 hours starvation to avoid fly death. Detailed methods for how FLIC operates can be found in Ro et al., 2014^39^. Fly feeding was measured for the first three hours in the FLIC and all FLICs were performed at 10 am local time. For each FLIC, half of the wells (n=6/FLIC) contained the fed group, and the other half contained the starved group of flies for direct comparison. 12-30 flies were measured for analysis of feeding. Any signal above 40 (a.u.) was considered a feeding event. Analysis of feeding events was performed using R.

### Triglyceride Measurement

Whole body TAG measurements were performed in accordance with previously published methods^95^. In Brief, whole flies (n=3) were used per biological replicate with 8-9 replicates used for each timepoint and treatment. Flies were collected for TAG for all timepoints and treatments measured for feeding behavior (see above). Data was normalized to TAG levels per number of flies. Significance was calculated using two-way ANOVA.

### Gene Expression

30 fly fat bodies (For Pect) or heads (for Dilps) of each genotype were dissected in RNAlater. Immediately after dissection, fat bodies were moved to tubes of 200uL RNAlater on ice. RNAlater was then removed, 30uL of trireagent and a scoop of beads were added, and fat bodies were homogenized using a bullet blender. RNA was then isolated using a Direct-zol RNA microprep kit following the manufacturer’s instructions. Isolated RNA was synthesized into cDNA using the BioRad iScript RT supermix for RT-qPCR and qPCR was performed using the BioRad ssoAdvanced SYBR green master mix. Primers used in the paper are as follows: Robl (endogenous control), forward: AGCGGTAGTGTCTGCCGTGT and reverse: CCAGCGTGGATTTGACCGGA; Pect, forward: CTGGAAAAGGCTAAGAAACTGGG and reverse: TCTTCAGTGACACAGTAGGGAG ; alpha-tubulin (endogenous control), forward: ATCGAACTTGTGGTCCAGACG and reverse: GGTGCCTGGAGGTGATTTGG ; Dilp2, forward: GCCTTGATGGACATGCTGA and reverse: CATAATCGAATAGGCCCAAGG; Dilp5, forward: GCTCCGAATCTCACCACATGAA and reverse: GGAAAAGGAACACGATTTGCG; Pect. Relative quantification of mRNA was performed using the comparative CT method and normalized to Robl mRNA expression. For each experiment, three biological replicates were used with 3 technical replicates used for qPCR.

### Statistical Methods

t-test or ANOVA were performed using PRISM. Boxplots and standard deviation calculations were either performed through prism or through R. R packages used in this paper included tidyverse, ggplot2, and ggthemes.

---

**Figure S1.**
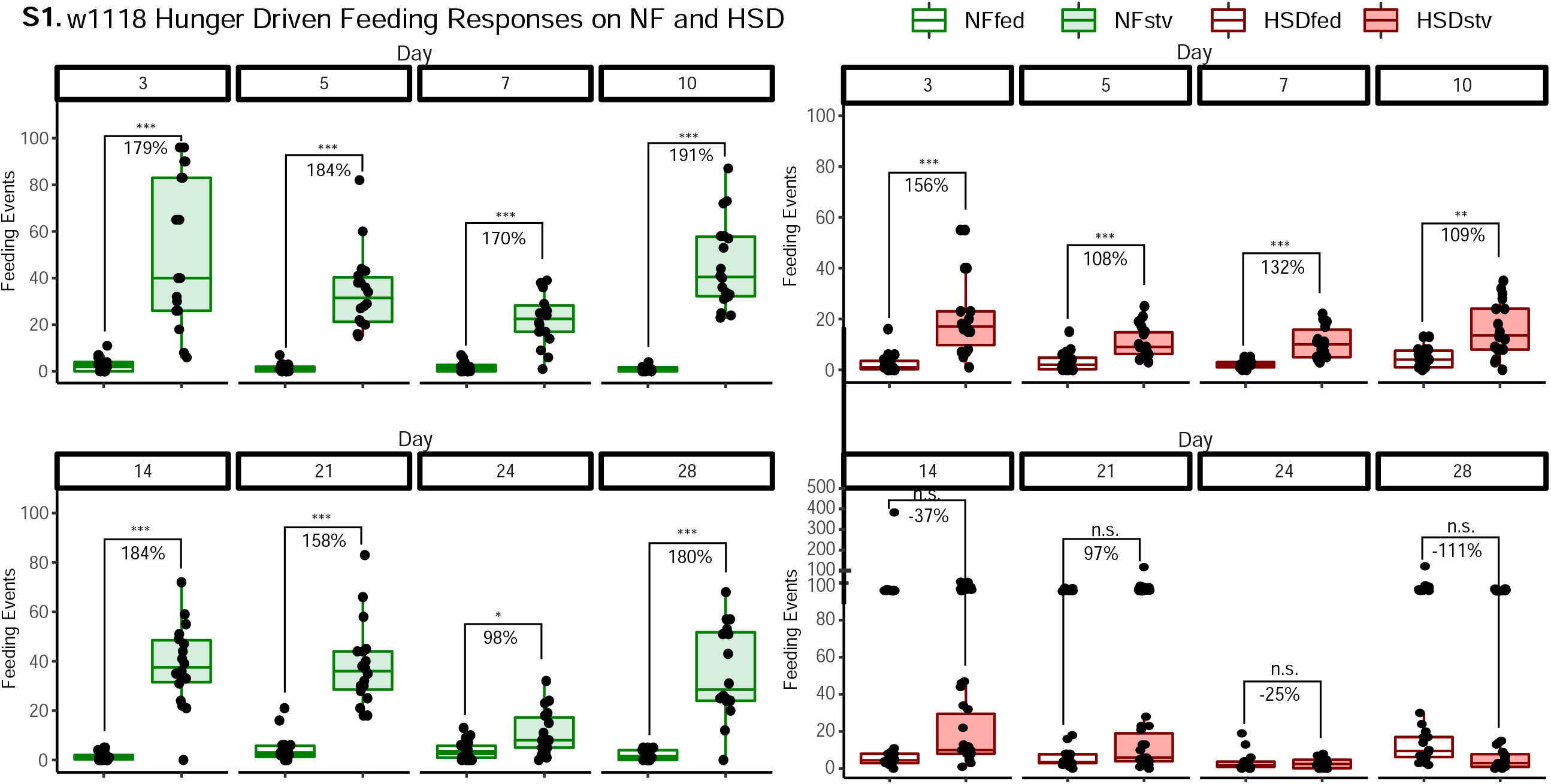
Flies on HSD show a progressive loss in hunger-driven Feeding. A, left) Average feeding events over time for NF flies that were either fed or starved for 16 hours prior to measurement (stv). Note that HDF is maintained throughout the experiment. A, right) Average feeding events over time for HSD-fed flies that were either fed or starved for 16 hours prior to measurement (stv). Note loss of HDF at day 14. Each dot denotes an individual fly. Two-Way ANOVA with Sidak post-test correction. Asterisks indicate significant changes with *p value <0.05, **p value <0.005, and ***p value <0.0005. Error bars= standard deviation.

**Figure S2.**
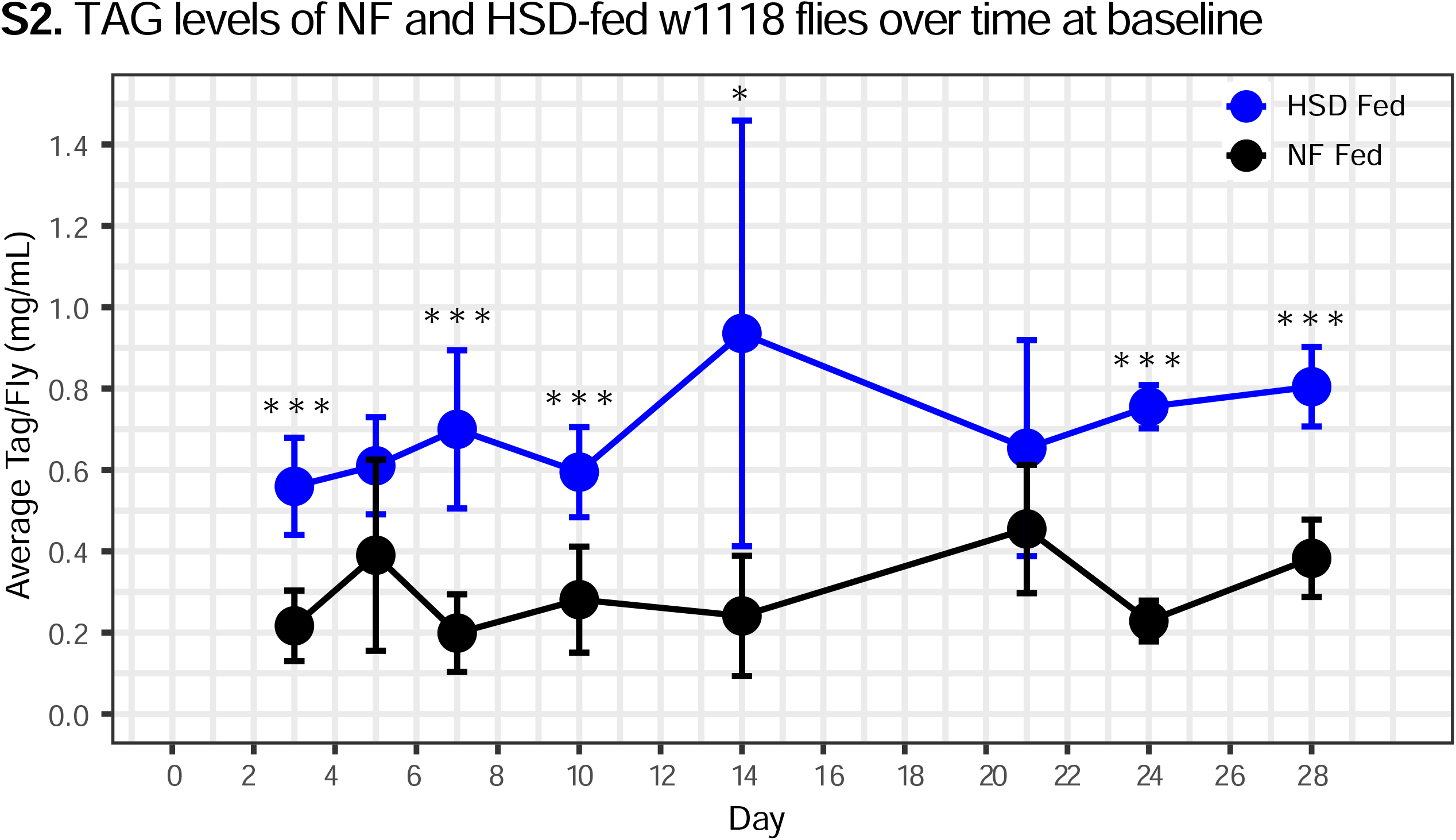
NF and HSD flies show similar rates of TAG breakdown following a starvation challenge. A) Average TAG/Fly levels in NF (black line) and HSD (blue line) groups over time. N= 9/group. 2-way ANOVA with Sidak post-test correction. Asterisks indicate significant changes with *p value <0.05, **p value <0.005, and ***p value <0.0005. Error bars= standard deviation.

**Figure S3.**
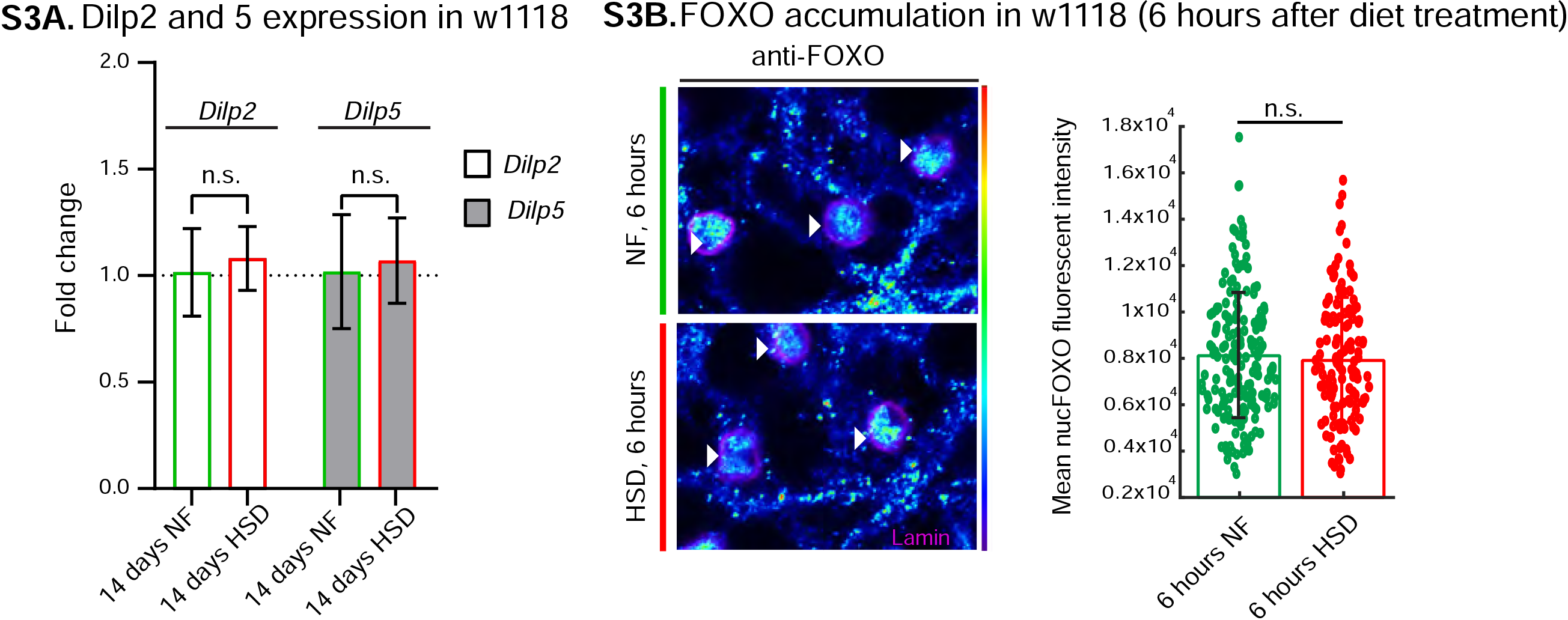
Acute HSD exposure does not affect FOXO nuclear localization. A) Mean fold change in Dilp 2 and Dilp 5 expression in w1118 flies fed either a NF or a HSD for 14 days. N= 3 technical replicates of cDNA collected from 30 flies/ treatment. T test with Welch’s correction showed no significant difference. B) Representative confocal images of nuclear FOXO accumulation in the fat bodies of NF (top panel) and HSD-fed flies for 6 hours (bottom panel). The nuclei are marked with anti-lamin (magenta). Arrowheads point to nuclei. B, Right) B, Right) Mean nuclear FOXO fluorescent intensity from z-stack summation projections of fat bodies from NF and HSD-fed flies. N=each circle represents a nucleus. Two-sided Wilcoxon rank-sum test. Error bars=standard deviation.

**Figure S4.**
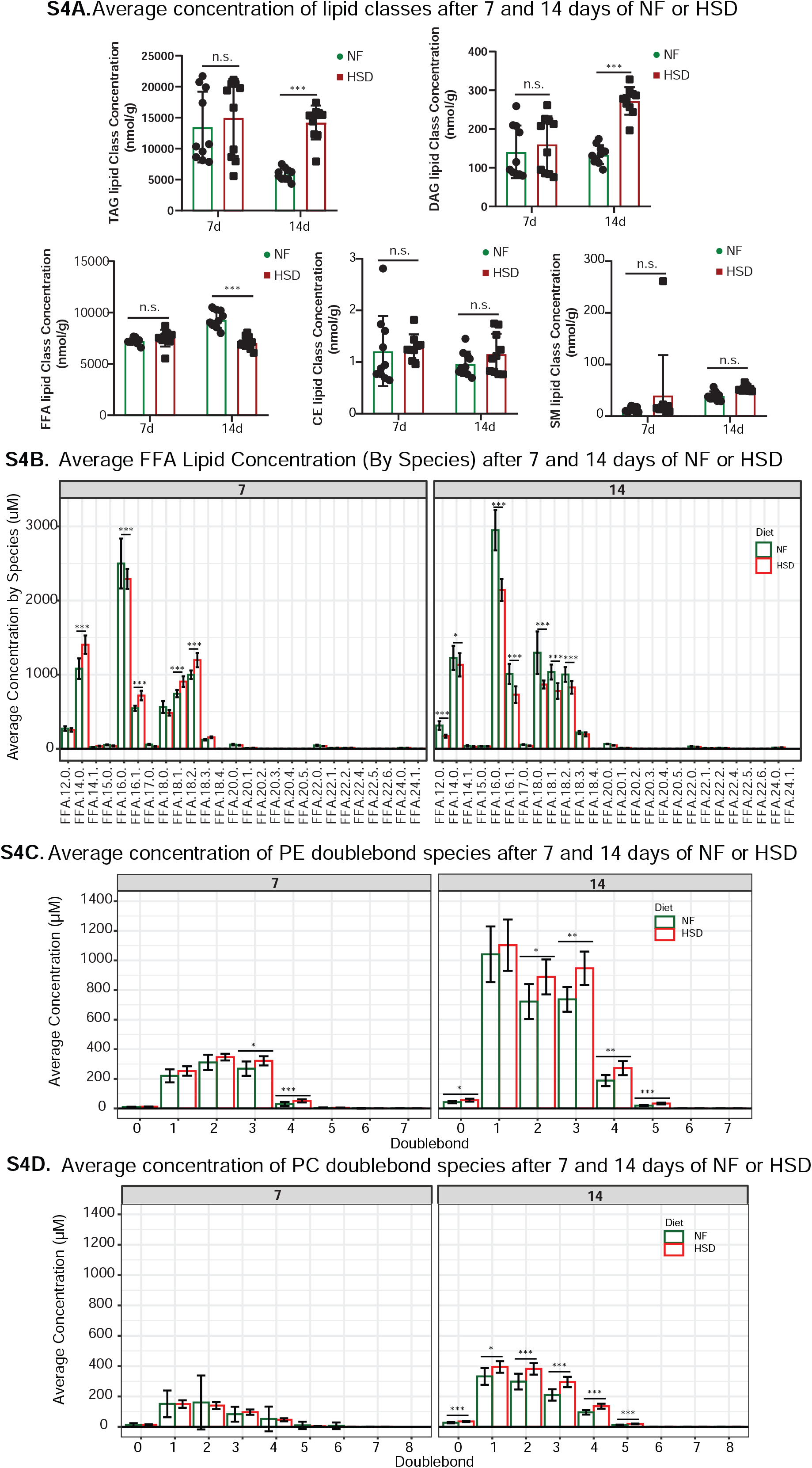
HSD-induced changes in the concentration of PE and PC double bond species. A) changes in TAG, DAG, FFA, CE, and SM concentration after 7 and 14 days of NF and HSD treatment. B) depicts changes in concentration of individual FFA species in HSD compared to NF at days 7 and 14. Lipid species were assessed for changes in concentration based on doublebond number for PE (C) and PC (D). Concentrations for species with the same double bond number were summed together and the difference between concentrations of HSD to NF was compared for days 7 and 14. n=10 flies/ condition. 2-way ANOVA with Holms Sidak correction. Asterisks indicate significant changes with *p value <0.05, **p value <0.005, and ***p value <0.0005. Error bars= standard deviation.

**Figure S5.**
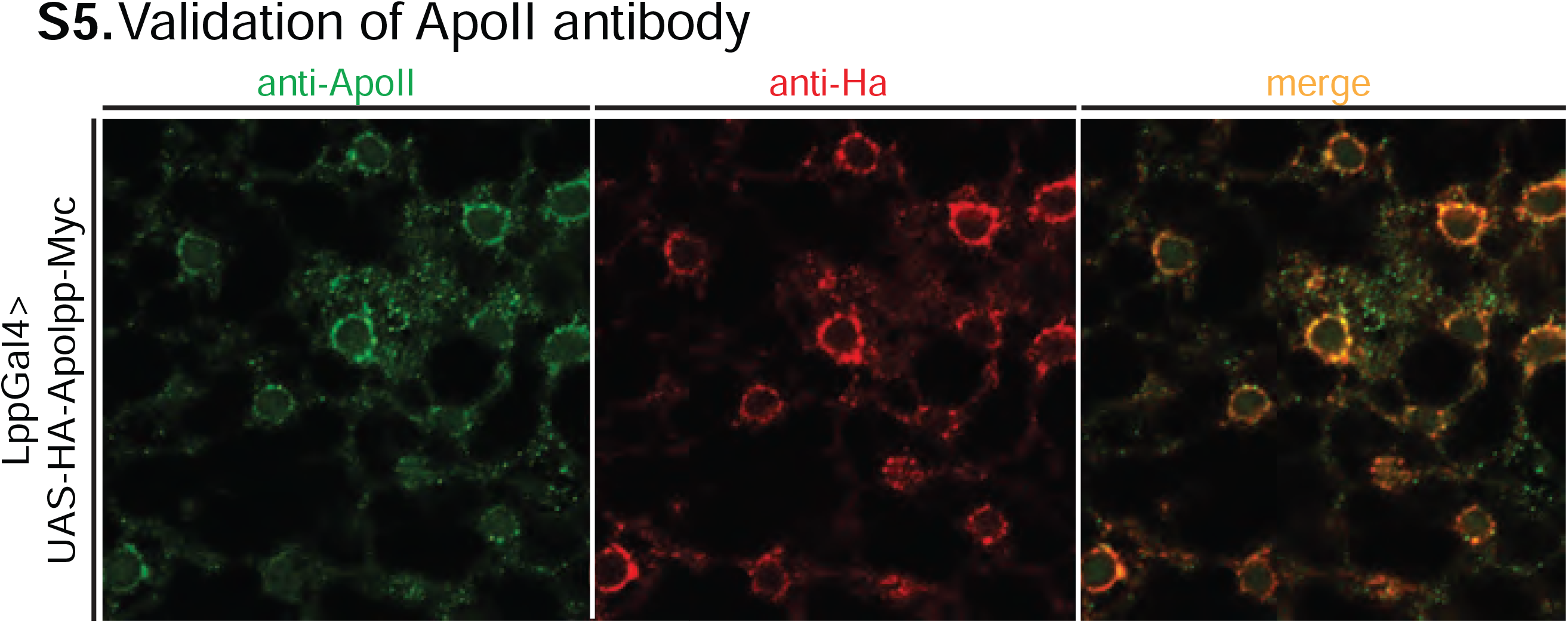
validation of ApoII antibody. Representative confocal images of LppGal4>UAS-HA-Apolpp-myc fly fat body stained for ApoII antibody (green), HA antibody (red), and a merge of the two images. Note colocalization of HA and ApoII.

**Figure S6.**
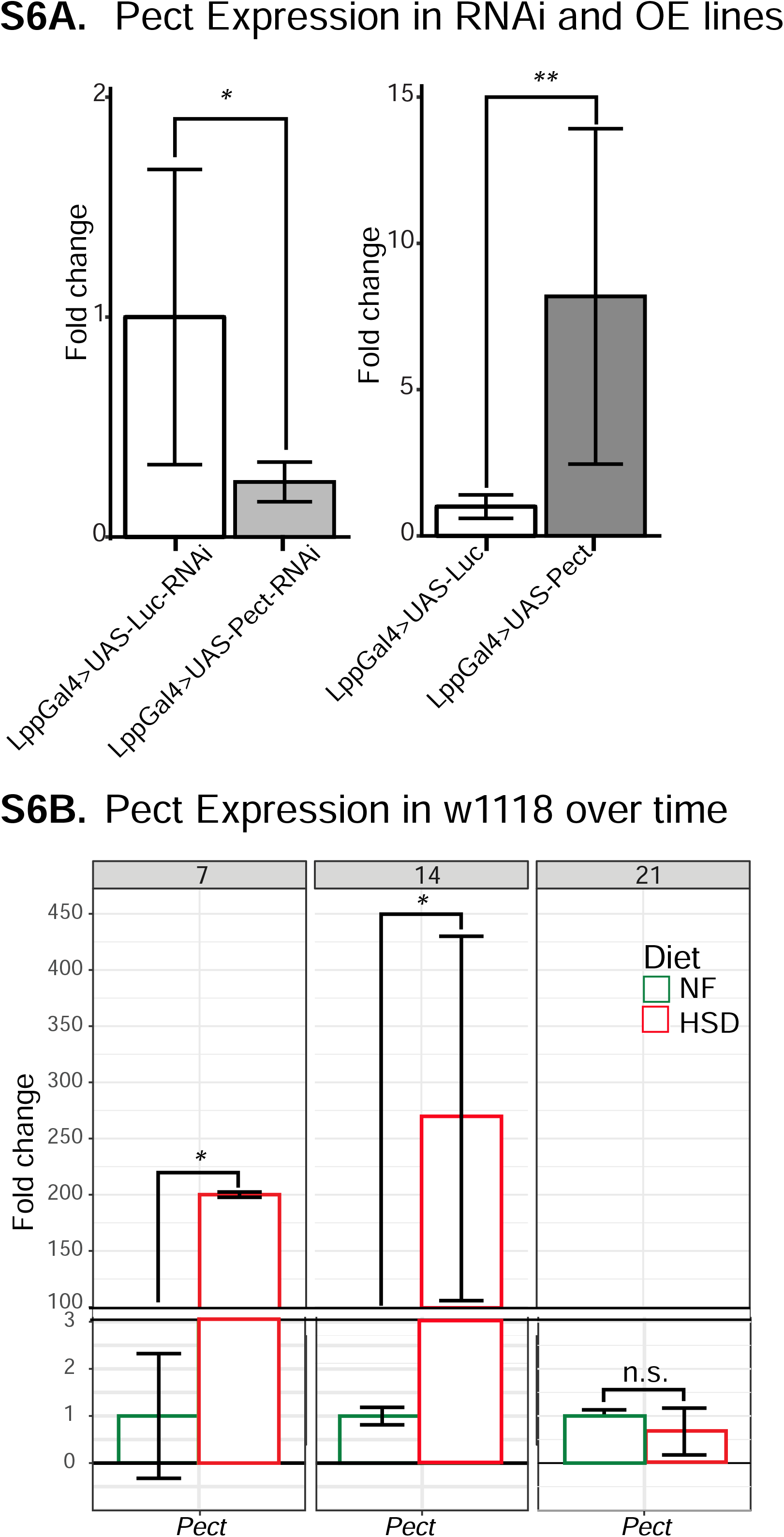
Pect mRNA expression. A) Mean fold change in Pect mRNA levels in the fat-specific (left) Pect knockdown flies and (right) Pect overexpression flies. Unpaired t-test with Welch’s correction B) Mean fold change in Pect mRNA levels of NF-fed and HSD-fed flies over time. Two-way ANOVA with Holms Sidak correction. N=3 technical replicates/group of cDNA collected from an N=30 flies/group. Asterisks indicate significant changes with *p value <0.05, **p value <0.005, and ***p value <0.0005. Error bars= standard deviation.

**Figure S7.**
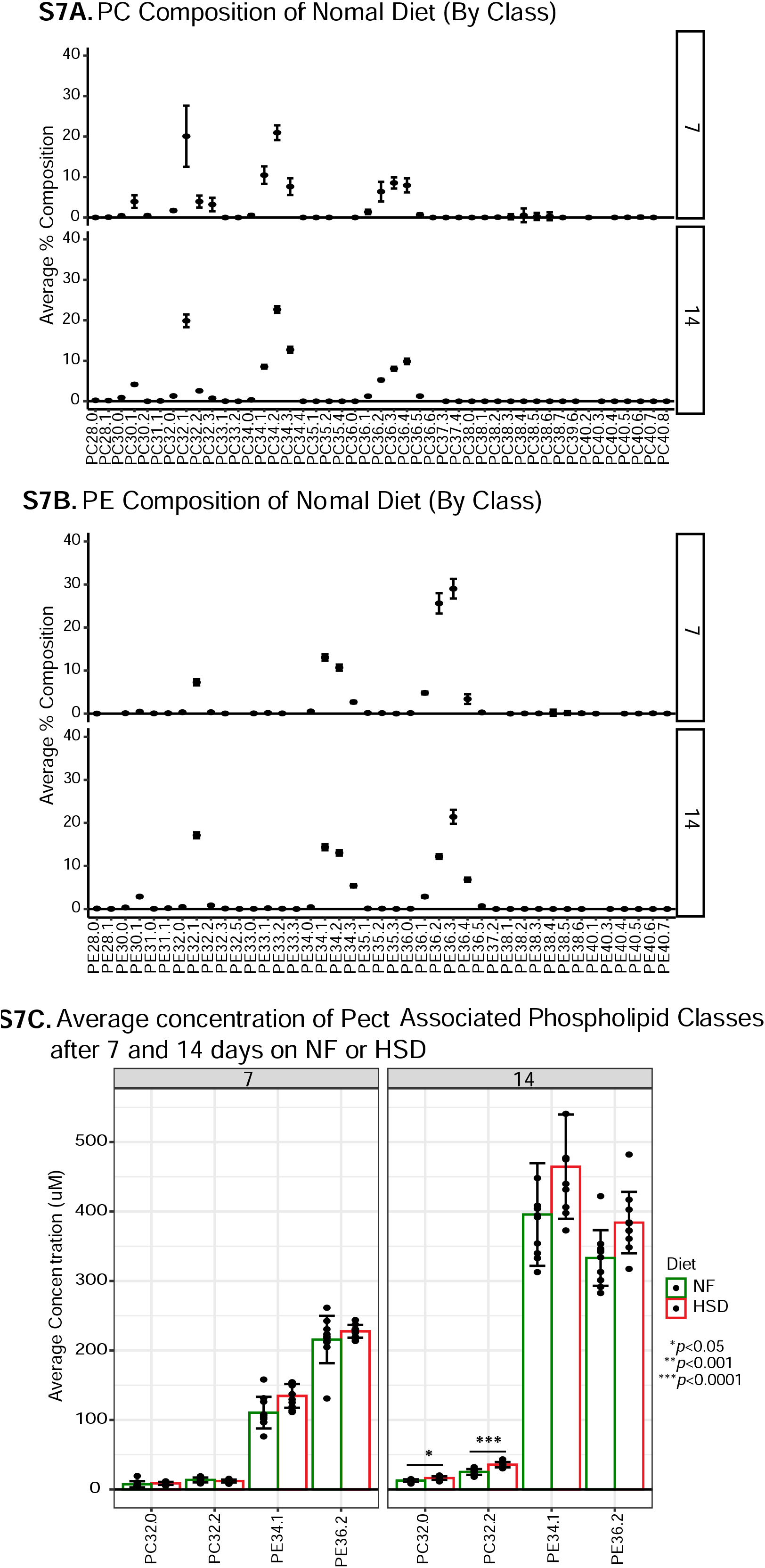
14 days of HSD causes an increase in Pect-associated phospholipid Classes. B) % lipid composition for all (A) PC and (B) PE classes in 7 NF flies overtime. Note that the overall composition does not change with age. The lipid composition was averaged amongst 10 biological replicates (n=10 flies/replicate). Error bars indicate standard deviation. C) Average concentration of PC and PE classes that are associated with Pect based on Tsai et al.^25^ Bars plot average concentration amongst 10 biological replicates (n=10 flies/replicate). Two-Way ANOVA with Holms Sidak correction. Asterisks indicate significant changes with *p value <0.05, **p value <0.005, and ***p value <0.0005. Error bars indicate standard deviation.

**Figure S8.**
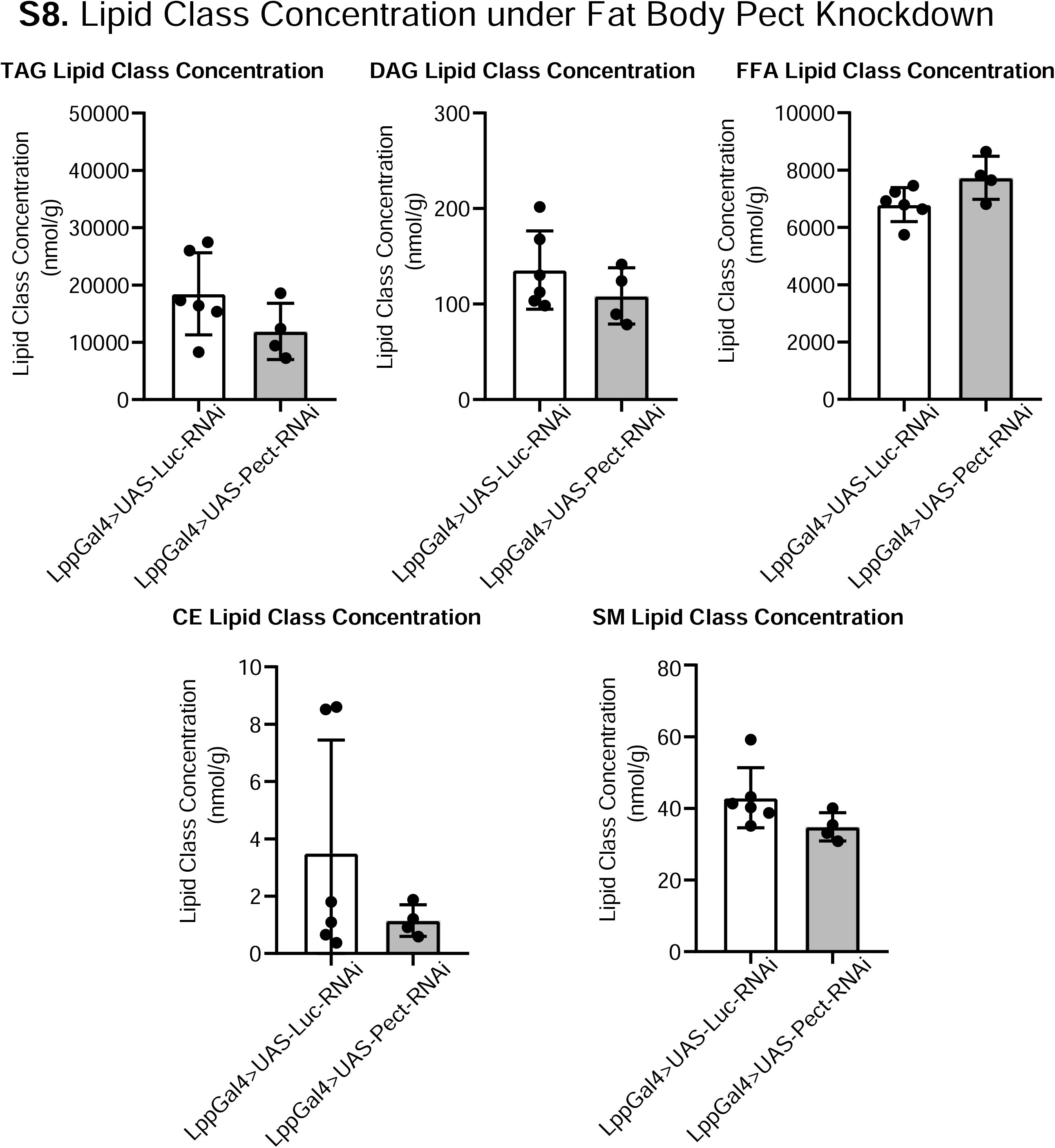
Additional lipid class responses to fat body Pect knockdown. Average concentrations of A) TAG, B) CE, C) DAG, D) SM, and E) FFA lipid classes in LppGal4>UAS-Pect-RNAi flies compared to control under 7 day NF conditions. Lipidomics was performed using a targeted quantitative lipidyzer (Sciex 5500 Lipidyzer). 4-6 independent biological replicates were used for each genotype, with n=10 flies composing 1 biological replicate. Unpaired t-test with Welch’s correction. Asterisks indicate significant changes with *p value <0.05, **p value <0.005, and ***p value <0.0005. Error bars= standard deviation.

**Figure S9.**
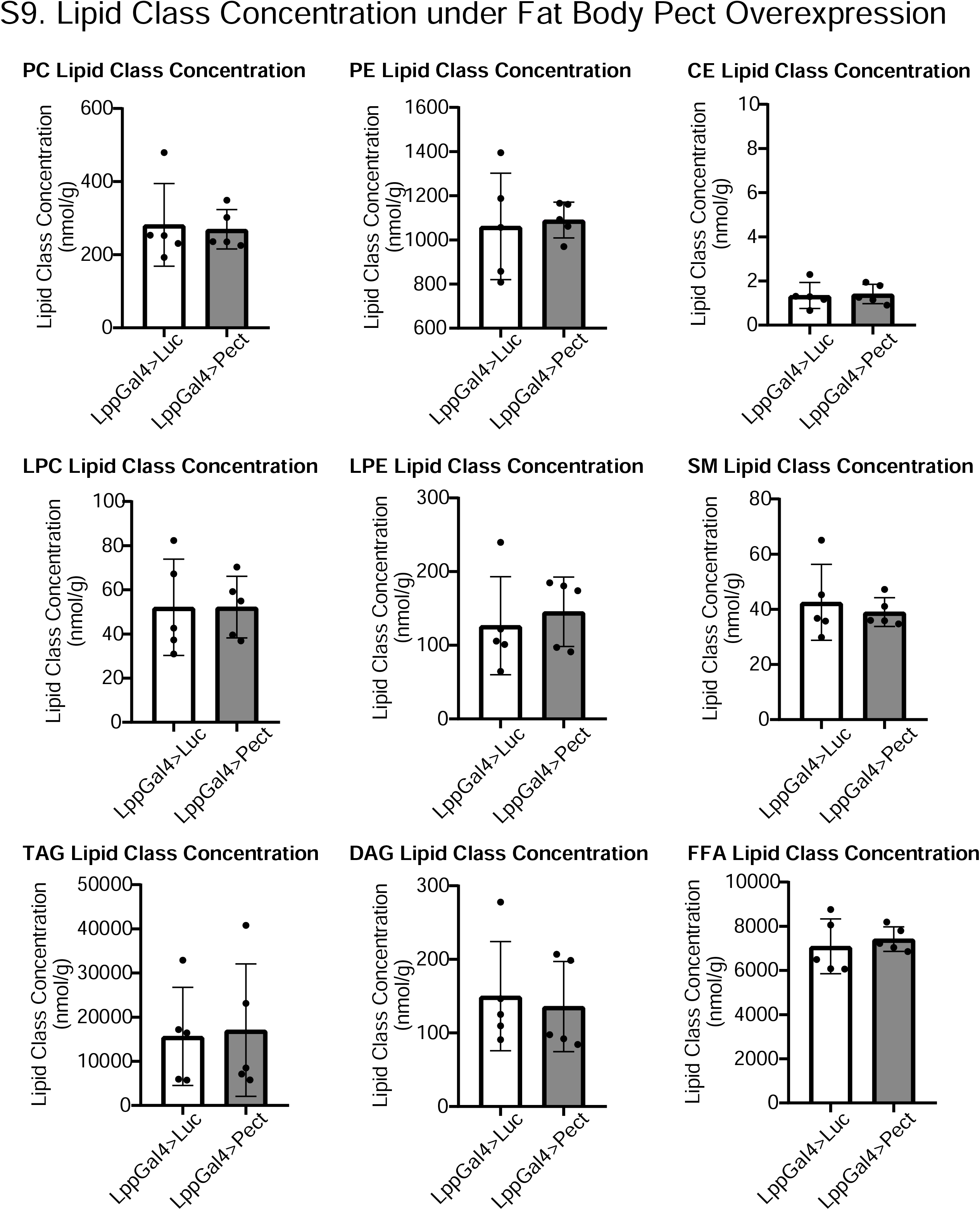
Lipidomic profile of fat body Pect overexpression flies. Average concentrations of A) PC, B) PE, C) CE, D) LPC, E) LPE, F) SM, G) TAG, H) DAG, and I) FFA lipid classes in LppGal4>UAS-Pect flies compared to control under 7 day NF conditions. Lipidomics was performed using a targeted quantitative lipidyzer (Sciex 5500 Lipidyzer). 4-6 independent biological replicates were used for each genotype, with n=10 flies composing 1 biological replicate. Unpaired t-test with Welch’s correction. Asterisks indicate significant changes with *p value <0.05, **p value <0.005, and ***p value <0.0005. Error bars= standard deviation.

## Full Revision

### 1) General Statements [optional]

> *This section is optional. Insert here any general statements you wish to make about the goal of the study or about the reviews.*

The goal of this study is to:

- Define how prolonged exposure to a high-sugar diet (HSD) regime alters both the lipid landscape and feeding behavior.
- Determine how changes in lipid classes within the adipose tissue regulates feeding behavior.

Key findings:

In this study, by taking an unbiased systems level and genetic approach, we reveal that phospholipid status of the fat tissue controls global satiety sensing.

Impact of Key findings:

By uncovering a critical role for adipose tissue phospholipid balance as a key regulator of organismal feeding, our work raises the possibility that the rate-limiting enzymes in phospholipid synthesis, including Pect, are potential targets for therapeutic interventions for obesity and feeding disorders.

Peer review comments:

This study has immensely benefited from the thoughtful peer-review of three reviewers. As per their recommendations, we have performed a major revision by performing additional experiments (see summary table below in next section) and strived to address the major concerns raised. Based on our reading, there were two major concerns that overlapped between all three reviewers raised. They are as follows:

- Does the genetic disruption of Pect in fly fat body alter phospholipid levels? Two reviewers (#2 and #3) recommended that we perform lipidomic analyses on adult flies with adipose tissue specific knockdown of Pect. For the revised version, we have completed this lipidomic experiment, and present results as a new main Figure 6, Supplemental S7 and S9.
- Is the dampened HSD induced hunger-driven feeding (HDF) behavior because of increased baseline feeding (#1 and #3)? In addition, reviewer #1, asked us whether HSD flies experience an energy-deficit? In other words, we were asked to uncouple whether what we observed was HSD-driven allostasis or indeed, as we had interpreted, that HSD dampened hunger-driven feeding response. Hence, they recommended that we:

i. Re-analyze our hunger-driven feeding datasets and present non-normalized data (also requested by Reviewer #3) and show baseline feeding behavior on HSD. To address this, we have completed this analysis and present our results in Figure 1B-D and S1.
ii. Determine whether the HSD fed flies display an energy deficit on starvation. To this end, we performed an assayed starvation-induced fat mobilization on HSD, results for this are now presented on Figure 1E-G and S2.

Conclusions after the revision:

First, it is important to note here that the additional experiments have not caused a significant revision of the major conclusions of the original version of our study. In fact, we hope that the revised version provides clarity and further substantiation to our original arguments.

- The lipidomics experiments on Pect fat-specific knock-down flies show that reducing Pect in fat-body causes a significant reduction in certain PE lipid species (PE 36.2 specifically-Figure 6B). This is consistent with a prior report on lipidomics of the Pect null allele by Tom Clandinin’s group (PMID: 30737130). Furthermore, we note that when Pect is knocked down in the fat body, there is a significant increase in two other classes of phospholipids LPC and LPE (Figure 6A). Together, this suggests that an imbalance in phospholipid composition in the absence of Pect activity in fat.
- The starvation-induced fat mobilization experiments show that despite being fed a prolonged HSD, adult flies sense starvation and effectively mobilize fat stores, at a level comparable to Normal food (NF) fed adult flies, suggesting that even despite HSD exposure, adult flies experience an energy deficit on starvation.
- In our non-normalized data, we find that the baseline feeding events are *not significantly altered between HSD and NF-fed flies (Figure 1D)*. This suggests that the effects we observe are not due to an increase in the “denominator”, but a dampening of hunger-driven feeding on HSD.

With regard to our original version, all three peer-reviewers found that the study was interesting, significant, important, and novel – Reviewer #1: “*The work is potentially novel and interesting*”; #2 : “*I find the study to be potentially very important - the authors combine a longitudinal study that would be difficult in any other model with the powerful genetic tools available in the fly. The conclusions are mostly convincing*”; #3: “*This manuscript demonstrates how fat body Pect levels affect HSD induced changes in hunger-driven feeding response. I agree with all the reviewers points; potentially very interesting*”. But had requested that we provide further substantiation and clarification.

We sincerely hope that the peer-reviewers find that our revised version with additional new experimental datasets, improved data visualization, and the presentation of non-normalized raw data points, makes this study clear, compelling, and well-substantiated.

### 2) Point-by-point description of the revisions

> *This section is mandatory. Please insert a point-by-point reply describing the revisions that were <u>already carried out and included</u> in the transferred manuscript*.

Below we summarize in Part A, the key experiments that were performed to address the major concerns. In Part B, we provide a point-point response to each reviewer with embedded datasets.

Part a:

We performed several new experiments, including:

1. To address the primary concern of Reviewer #1 regarding whether the HSD flies have a similar energy deficit to Normal food (NF) fed flies, we performed analysis of stored neutral fat Triacylglycerol (TAG) reserves and how HSD fed flies mobilized fat stores on starvation. We present these results in Figure 1E-G, S2. These results show that HSD-flies despite accumulating more TAG (S2), breakdown a similar amount of fat reserves as NF-fed flies on starvation at any time-point (Figure 1E-G). This suggests that HSD-fed flies do sense and respond to energy deficit.
2. To address concerns of reviewer #2 and #3 on whether Pect genetic manipulation affects specific phospholipid classes, we performed lipidomic analyses.

The table below summarizes the new 3 new figures and 4 supplemental figures (blue text are all new figure numbers and figure panels) and three new Supplementary files as per reviewer’s request.

| Figure # | Main point | New datasets in revision | Companion Supplement |
| --- | --- | --- | --- |
| 1 | HSD alters feeding behavior, but flies still breakdown TAG on starvation. | TAG storage and breakdown over longitudinal HSD shows that HSD and NF fed flies show similar levels of TAG breakdown on starvation, despite consistently elevated TAG on HSD. This supports the idea that flies do sense starvation even on HSD, but there is a uncoupling of the feeding behavior after Day 14. Revised the data representation of Figure 1 to show non-normalized data over time. S1 and S2 companions are new in the revision. Panels 1D to 1E are new for the revision. | S1- Raw data of feeding events plotted.<br>S2 Elevated TAG at all time points. |
| 2 | HSD causes insulin resistance | S3A added to show that insulin transcript levels remain the same in response to reviewer #3's concerns. | S3 |
| 3 | Phospholipid concentration raw data from lipidomic on Day 7 and Day 14 HSD suggest that PC, PE levels are increased on Day 14 HSD. | Figure 3 revamped to show new data visualization and non-normalized raw data to address Reviewer #2's major concerns. S4A and S4B added. In addition Supplementary File 1 and 2 provided with raw lipidomics data as per reviewer #2's request. | S4.<br>S4A- non normalized raw data of all other lipid classes on HSD.<br>S4B- fatty acid species data on Day 14 added as per request of rev.#2. |
| 4 | HSD regulate Apo-I levels in the IPCs and phenocopies Pect KD. | Added Figure 4A to show that HSD phenocopies Pect-KD in terms of delivery to brain | S5 showing the validation of the Apo-I antibody.<br>S6 validation of Pect KD and over-expression and Pect mRNA levels dysregulation on HSD. |
| 5 | Pect RNAi is insulin resistant | N/A | N/A |
| 6 | Pect knockdown shows significant increase in LPC and LPE, and a non-significant reduction in PC, PE levels. Specifically, the PE lipid class PE36.2 is downregulated. | Fig 6, S7, S9 are completely new based on reviewer #2 and #3 requests. In addition Supplementary File 3 provided with raw lipidomics data as per reviewer #2's request | S7, S8, S9#.<br>S7- new Pect KD other classes<br>S8- new PE classes for day 14 and Pect associated classes.<br>S9- Pect OE lipidomics |
| 7 | Pisd and Pect activity in adipocytes are required for hunger-driven feeding behavior in normal diets | Pisd RNAi data was moved from supplement to main figure. | N/A |

*Note on revised text:* We have revised text not only in the results section, but also as per reviewer #2’s recommendation, we have revamped our introduction and discussion as well. Since the manuscript has been significantly revised to include a main figure 6, fully altered Figure 1 and 3, multiple new supplemental figures*, the changes in text are extensive. Hence, they are unmarked in the main text.* Nonetheless, we hope that the reviewers will be able to evaluate these changes, as we have provided the specific locations in text and embed key figures in the point-point response below.

**<u>Part B:</u>** Point-Point responses to reviewer comments.

Reviewer #1 comments in Blue, author response in black.

Reviewer #1 (Evidence, reproducibility and clarity (Required)):

In this manuscript, Kelly et al. show that the difference between the feeding behavior of fed and starved flies (hunger-driven feeding; HDF) is absent in animals fed a high-sugar diet (HSD) for two weeks or more. The disappearance of HDF with HSD coincides with changes in phospholipid profiles caused by HSD. Furthermore, RNAi-mediated downregulation of Pect in the fat body-a key enzyme in the PE biosynthesis pathway-phenocopies physiological effects of HSD. Moreover, downregulation or overexpression in the fat body abolishes or induces HDF, respectively, abolishes or induces HDF, respectively, independent of HSD treatment.

Overall, the manuscript is well-written and the phenotypes are clear. However, I have major concerns regarding the authors’ interpretation of the data and their conclusion. Most importantly, while it is clear that the authors’ high-sugar dietary treatment affects feeding behavior and physiology, I am not convinced that the changes can be considered “hunger-driven”-which is central to the main point of the manuscript. Therefore, it is my recommendation that the authors substantially revise the manuscript by either showing additional/re-analyzed data that rule out alternative hypotheses, or rewriting the manuscript keeping alternative interpretations in mind.

We are thankful to this reviewer for their thoughtful critique, and constructive and specific suggestions on how we can redress these concerns. We have taken on board the concerns of this reviewer regarding our interpretation of whether the changes in feeding behavior can be considered hunger-driven or not. Based on their advice, we have made significant changes by addressing: i) does HSD increased baseline feeding-we now show non-normalized raw data and data supports conclusion that baseline feeding is not higher; ii) whether HSD-fed flies can sense an energy deficit at levels similar to NF fed flies-we show that HSD flies sense energy deficit. We have provided detailed response below, and we hope the reviewer finds the additional datasets and re-analyzed data are consistent with the interpretation that prolonged HSD dampens starvation induced feeding. In addition to this key concern this reviewer has made a many other salient points that we have addressed with additional data or by clarifying the text.

Major comments:

1) The data do not sufficiently show that the long-term HSD regime disrupts “hunger-sensing.” The manuscript should address alternative hypotheses by showing raw instead of normalized data, rewriting the manuscript with a new central conclusion, or running additional experiments that actually show a defect in hunger-driven response.

a. The main results that the authors rely on for the argument is that the ratio of feeding events that the starved and non-starved flies eat is different between the groups fed normal or HSD. However, because the authors only show normalized data (normalized to non-starved flies; Fig. 1), it is difficult to tell whether the change is due to a chronically increased feeding in non-starved HSD flies-maybe in perpetual hunger-like allostasis-or dampened starvation response. Indeed, the data shown in Fig S1 show that flies fed HSD for as short as 5 days show more frequent feeding events compared to age-matched controls fed normal food. It is possible that because the HSD-fed flies eat more than NF-fed flies, even without being starved, the ratio of starved/non-starved feeding is lower in the HSD-fed group-due to changes in the denominator, rather than the numerator.

We have taken onboard this concern regarding presenting only normalized data, and that clouded the interpretation and left open other possibilities. In the completely revised figure 1 and S1. We now show non-normalized data, as a function of time. First we note that HSD-fed flies, do not show higher baseline feeding that NF fed flies, except on Day 10 of HSD, when there is a modest but significant elevation (Figure 1D).

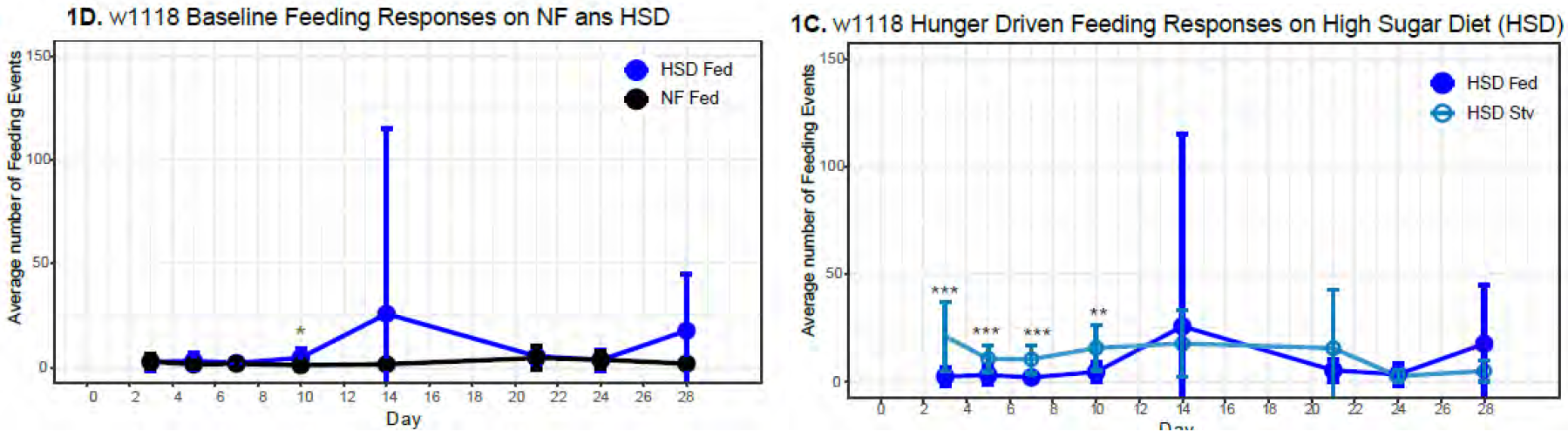

Nonetheless, on Day 10 HSD, flies still display increased hunger-driven feeding HDF (Figure 1C), it is only after Day 14 HSD that HSD dampens the starvation induced feeding.

b. It is also possible that the HSD-fed flies are simply not in as big an energy deficit physiologically, due to the increased fat deposits they’ve accumulated (as the authors show later in the manuscript). It may take longer for the fat HSD flies to reach substantial energy deficiency than the NF flies, but they still may eventually be able to appropriately respond to hunger, just like NF flies. In such case, it would be a misnomer to call this behavioral change a ‘defect in hunger-driven feeding behavior.’ Maybe an experiment with a dose-response curve of “hunger driven feeding response” as a function of duration of starvation would help?

Prompted by this reviewers question, we asked whether HSD fed flies, that have a higher baseline neutral fat store (Triacylglycerol-TAG) level, and if HSD-fed flies can sense energy deficit. For this, we revisited the longitudinal assays for neutral fat triacylglycerol (TAG) storage that our lab had generated, along with the HSD-HDF studies. We now present this evidence as Figure 1E-1G and Figure S2. Overall, our experiments point to the idea that adult flies fed HSD, are able to sense and mobilize TAG stores effectively throughout the 28-day time point that we analysed.

First as shown in Figure S2, flies fed HSD display an increase in TAG levels. But it is to be noted that while TAG stores increase, the increase is not linear with time. This suggests that adult flies exposed to HSD store excess energy as TAG, but the increased TAG stores stay within a certain range despite the length of HSD exposure. This suggests that adult flies on HSD still display TAG homeostasis.

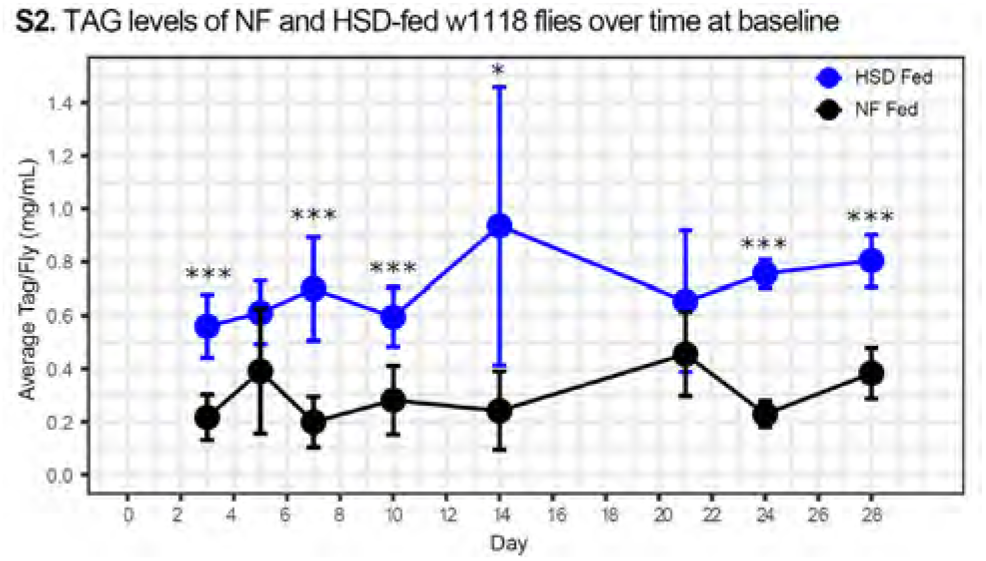

Next, to directly address the reviewers point about HSD fed flies not sensing an energy deficit, we subject HSD-fed flies to an overnight starvation, same regime as used in the overnight feeding experiments, and asked whether they mobilize TAG. We noted that flies exposed to HSD breakdown TAG throughout the 28-day exposure at statistically significant levels for Day 3-Day 28, except on 14 and 21 days (Figure 1F). While there is TAG mobilization on Day 14 and 21, the difference is not statistically significant. Nonetheless, we note the same levels TAG breakdown for normal lab food (NF) fed flies on Day 14 and 21 (Figure 1E). Overall, HSD fed flies sense and display energy deficit, as measured by TAG store mobilization, throughout the 28 days of HSD exposure, at levels comparable to NF-fed flies (Figure 1G).

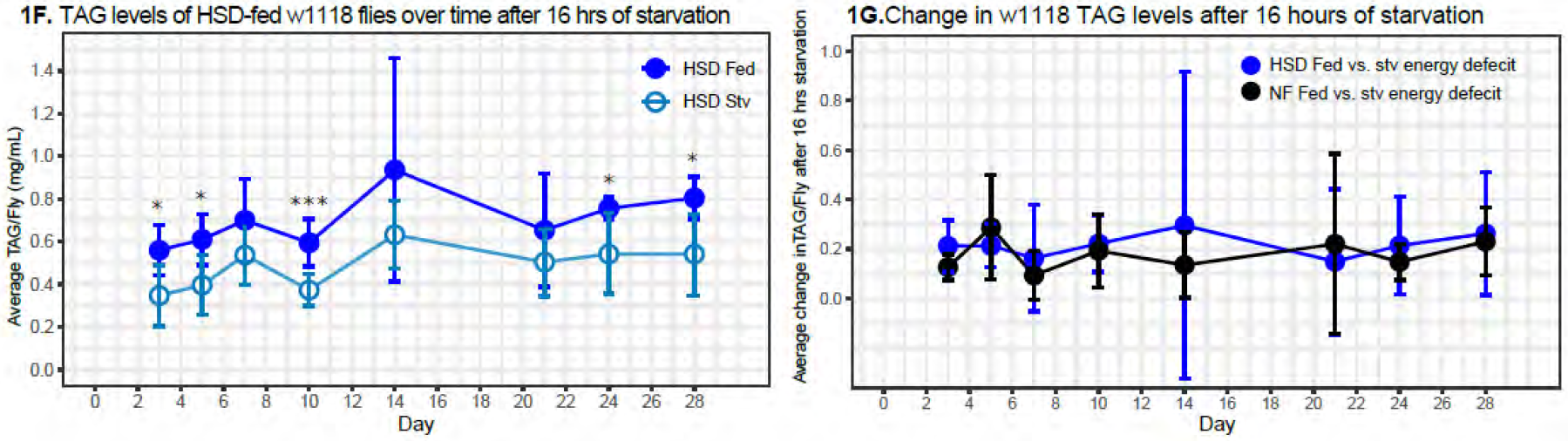

Taken together, these results suggest that while HSD-fed flies experience an energy deficit on starvation, at levels comparable to NF-fed flies, throughout the 28-day time point assayed. But, their starvation driven feeding-response is dampened by Day 14 and by Day 28, the HSD-fed flies display more feeding events than HSD starved flies. These results are consistent with the interpretation that in HSD-fed flies the starvation-induced feeding behavior becomes desynchronized from the starvation induced TAG-mobilization, suggesting that there is an absence of hunger-driven feeding.

2) How can you be sure that lower Dilp5 immunofluorescence is indicative of increased Dilp5 secretion? Wouldn’t decreased production of dilp5 also have the same results?

It has been shown previously in HSD fed larvae are hyperinsulinemic, i.e., they have 55% increase in circulating Dilp2 (PMID: 22567167). Additionally, we have shown that ectopic activation of the insulin-producing neurons by expressing TRPA1, an ion channel that activates neurons, reduces Dilp5 accumulation without a change in Dilp5 mRNA levels (PMID: 32976758), suggesting that reduced Dilp5 accumulation, without alterations to mRNA levels is a proxy for increased secretion. Now, in response to this concern, in the revised manuscript, we have added qPCR data of Dilp2 and 5 (Figure S3A), which show no difference in expression levels after 14 days on HSD. Therefore, there is no dip in Dilp5 mRNA production. Given that Dilp2 and Dilp5 mRNA levels remain the same, but we see reduced Dilp5 accumulation, we interpret this to mean that Dilp5 secretion is increased.

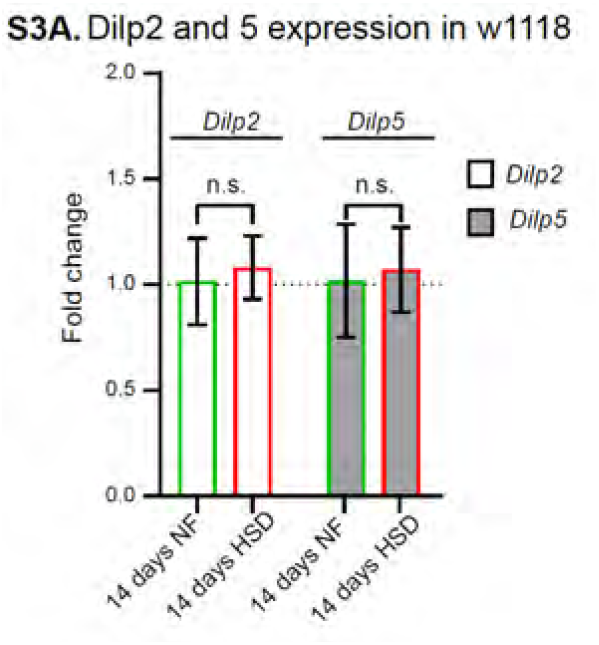

a. Also, the authors should state in the main text that it is Dilp5, not just any Dilp.

Thanks for this suggestion and we have fixed this and referred to Dilp5 specifically throughout the text in the results section.

3) Data presentation:

a. Sometimes the data are normalized to NF (Fig 4B-C), sometimes not (ex. Fig 4A, S4C). Unless there is a specific rationale for the data transformation, it would be more appropriate to show untransformed data (ex. Fig 4A, S4C), especially as the authors use two-way ANOVA to determine significance. Only showing the differences implies comparison against a hypothetical mean (i.e. μ0=0), not between two group means.

We thank the reviewers for bringing this issue to our attention. We updated all the figures to show untransformed data in the revised manuscript.

b. Some figures show both individual data points and summary statistics (mean, SD, … ex. Fig 2A)-which I believe is ideal-but some show only one or the other (ex. Fig 2B, no summary statistics; Fig. 3, no data points. The manuscript would read more convincing if data visualization is consistent across figures.

We thank the reviewers for their feedback. We have made changes to all the figures in the revised manuscript to improve visual consistency.

Minor comments:

1) High sugar diet: what is the actual sugar concentration in the NF v. HSD diets? The authors write that the HSD diet contains “30% more sugar” than the NF, but providing the final sugar concentrations-sucrose or others-would be informative for other scientists studying the effect of high sugar diets.

We thank the reviewer for their suggestion and now we have updated the methods to include this sentence. *“After 7 days, flies were either maintained on normal diet or moved to a high sugar diet (HSD), composed of the same composition as normal diet but with an additional 300g of sucrose per liter”*.

a. Additionally, the definition of HSD is inconsistent. Main text (Page 5, line 17) states that their HSD is “60% more sugar than normal media,” whereas the figure legend (Fig 1) and the Methods state that the HSD contains “30% more sugar.”

We apologize for this egregious typo in the figure legend! We have now fixed this to say 30% HSD. Only 30% HSD was used throughout this study.

2) Starvation medium: please provide justification for why the authors used 1% sucrose/agar for starvation medium, instead of plain agar/water that most labs use. At least clarify and provide a reference for the claim that the 1% sucrose/agar “is a minimal food media to elicit a starvation response.”

We are very grateful for this reviewer identifying this this methods description error and bring it to our attention. **<u>We used 0% sucrose agar for overnight starvation in this study</u>** as most labs do. The error occurred because we were using another manuscript from the lab to help draft the methods section (PMID: 29017032). In that study, where we assayed the effect of *chronic starvation* our lab used: “*1% sucrose agar for 5 days at 25C”*. However, in this current study, because we are testing ***acute effects of overnight starvation***, we are using 0% sucrose agar.

3) Pect mRNA level is higher with HSD. This is surprising because not only, as authors mention, is increased PC32.2 with HSD suggests lower Pect activity, but also because Pect RNAi phenocopies long-term HSD in HDF behavior, lipid morphology, FOXO accumulation in fat body. The authors speculate that the data “likely shown an upregulation in an attempt to mediate the Pect dysregulation occurring at the protein level.” If that were true, a western blot may be informative. Zhao and Wang (2020, PLoS Genetics) generated a Pect antibody that seems compatible with western blot applications. That being said, I don’t think such data is critical for the manuscript. I mention this simply as a suggestion for the authors. a. page 8, line 22-23, did you mean to write “Given how PC32.2 is elevated after 14 days of exposure to HSD, we assumed that Pect levels would be low for flies under HSD,” not “high?” Otherwise the subsequent 2 sentences don’t make sense.

We agree that the most confusing aspect of the study was that Pect mRNA levels being very high on Day 14 HSD, but nonetheless the effects of Pect-KD phenocopied HSD. To resolve this, we have now performed lipidomic analyses on whole adult flies, when Pect is knocked-down (KD) by RNAi in the fat tissue. We now present a new dataset in Figure 6. Two striking changes occur. They are:

i. Pect-KD shows increase in the phospholipid classes LPC and LPE (Figure 6A). In contrast, LPE is significantly downregulated on HSD Day 14 (Figure 3).
ii. Pect-KD shows a significant reduction in specific class of PE 36.2 (Figure 6B). Our data regarding increase in PE 36.2 agree with a previous lipidomic analyses of Pect mutant retina (PMID: 30737130). In contrast, PE 36.2 trends upwards on 14 day HSD (Figure S7C) though not significantly.

On 14-day HSD consistent with extreme upregulation of Pect mRNA fed flies (Figure S6A; Pect mRNA 200-250 fold), PE trends upwards on 14-day HSD (Figure 3) and PE 36.2 trends higher (Figure S7C). We note that on the surface of it PE and LPE *per se* are contrasting between 14-day HSD lipidome and fat-specifc Pect-KD. But there is a significant commonality that under both states there is an imbalance of phospholipids classes PE and LPE. Hence, we propose that maintaining the compositional balance of phospholipid classes PE and LPE is critical to hunger-driven feeding and insulin sensitivity. Hence, either increase or decrease, of these key phospholipid species, may lead to abnormal hunger-driven feeding.

We agree that a western blot would be informative as well, but we were unable to obtain the reagent from Dr. Wang’s group, precluding us from performing this request. See email snapshot.

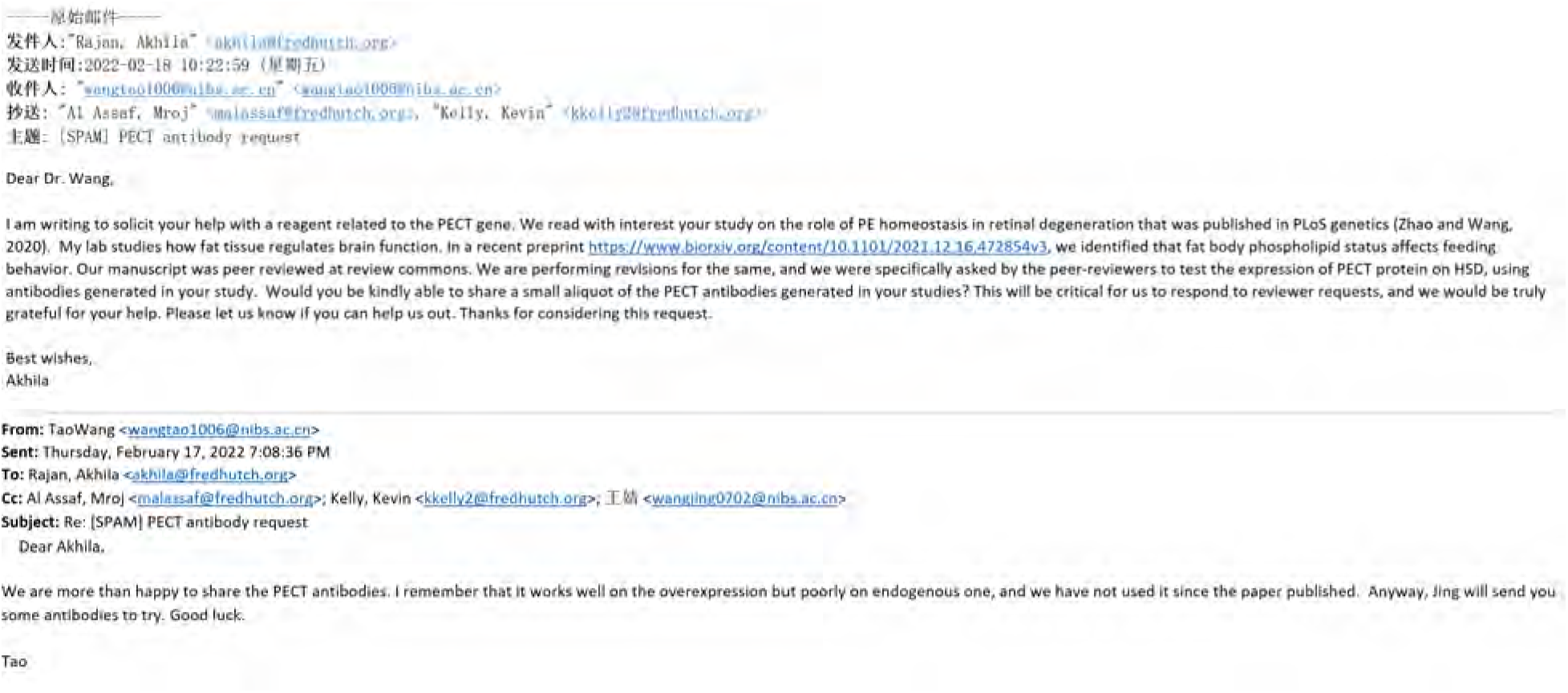

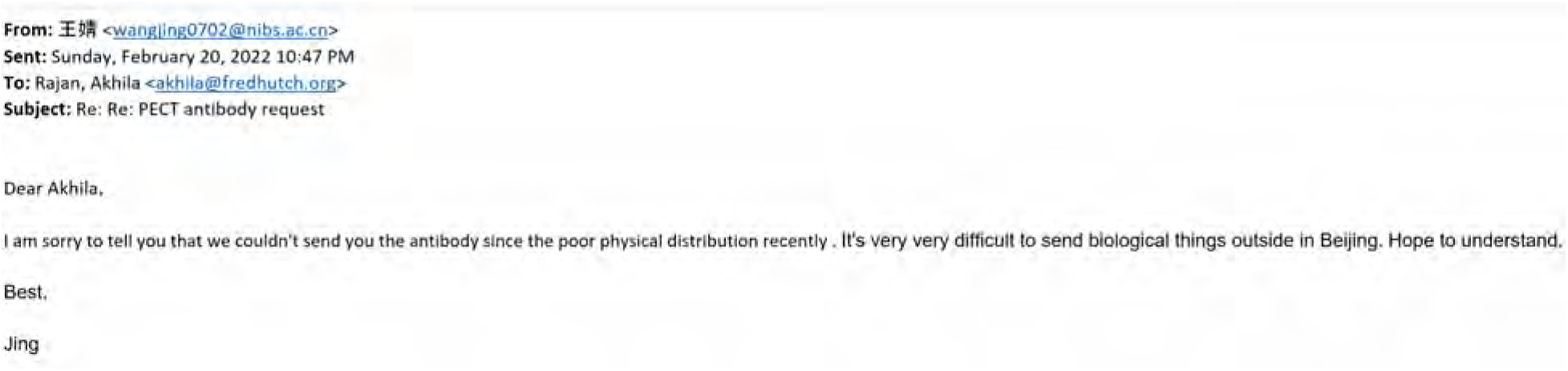

To ensure that we appropriately discuss and clarify this issue, we have now included a section in the discussion - Page 14 Lines 26-34-under the subtitle “*The implications of relationship between Pect levels and HSD*”. We have pasted an excerpt from that subsection below for this reviewers assessment.

> *“Also, we note that over-expression of Pect cDNA in the fat-body does not alter phospholipid balance (Figure S9) and indeed improves HDF on HSD (Figure 7B). While this may appear inconsistent, it is critical to note that over-expression of Pect cDNA using UAS/Gal4 only increases Pect mRNA expression by 7-fold (Figure S6A), whereas HSD causes its upregulation by 250-fold (Figure S6B). Hence, we speculate that an increased ‘basal’ level of Pect such as by that provided by a cDNA over-expression in fat, may be protective to the negative effects of HSD (Figure 7B) without affecting overall phospholipid levels (Figure S9), but extreme upregulation Pect on HSD affects the PE and LPE balance (Figure 3).”*

Reviewer #1 (Significance (Required)):

The work is potentially novel and interesting, but at this stage it’s difficult to interpret what the phenotype signifies. Although the manuscript could be revised simply by modifying the text, experimentally addressing the concerns would significantly improve the work.

In sum, we hope we have addressed the key concern for Reviewer #1 as to whether the behavior we report here is indeed a dampening of starvation-induced feeding, or an effect of increase in baseline feeding. We hope that by reviewing our non-normalized data, they can appreciate that it is the former. Also, we hope that Reviewer #1 appreciates that we have strived to address the concerns by additional experiments, to clarify our findings and improve the impact of the work.

**<u>Reviewer #2 (Evidence, reproducibility and clarity (Required)):</u>**

This intriguing manuscript by Kelly and colleagues uses the fruit fly Drosophila melanogaster as a model to understand how diet-induced obesity alters the feeding response over time. In particular, the authors findings indicate that chronic exposure to a high-sugar diet significantly alters the starvation-induced feeding response. These behavioral studies are complemented by a lipidomics approach that reveals how a chronic high sugar affects many lipid species, including phospholipids. The authors then pursue mechanistic studies that indicate phospholipid metabolism within the fat body appears to remotely affect insulin secretion from the insulin producing cells. Moreover, the changes in phospholipid abundance are associated with changes in insulin-signaling, including increased insulin secretion from the IPCs and elevated levels of FOXO within the nucleus.

I find the study to be potentially very important - the authors combine a longitudinal study that would be difficult in any other model with the powerful genetic tools available in the fly. The conclusions are mostly convincing, but a few follow-up experiments are required:

We are grateful for the reviewers constructive, detail-oriented, and balanced feedback, and their recognition of the value of this study. Now, we have performed additional experiments to address the key concerns raised by all reviewers. We hope that on reading the revised version of our study, that the reviewer continues to feel positive about the message of this study and its potential impact.

1. The key conclusions from the manuscript assume that manipulation of Pect expression levels alters phosphatidylethanolamine (PE) levels. However, the authors make no attempt to verify that the genetic experiments described herein actually affect PE levels. At a minimum, changes in PE levels should be verified for the Pect knockdown and overexpression lines. Similarly, there is no evidence that manipulation of either EAS or Pcyt2 induces the expected metabolic effects. I’m not asking that the longitudinal feeding experiments be repeated, simply that the authors measure the relevant lipid species, preferably with a targeted LC-MS approach.

Prompted by this reviewer, we performed targeted LC-MS on whole adult flies, on normal diet, to assess lipid levels for fat-specific Pect-KD and overexpression. We decided to focus on Pect, as its knock-down even on normal diet causes a dampened hunger-driven feeding behavior (Figure 7A) and phenocopied a 14-day HSD feeding phenotype.

We now present a new dataset in Figure 6. Two striking changes occur: They are:

i. Pect-KD shows a significant reduction in specific class of PE 36.2 (Figure 6B). Our data regarding decrease in PE 36.2 agree with a previous lipidomic analyses of Pect mutant retina (PMID: 30737130). It is to be noted that though overall levels of all PE species trend downwards, like the Clandinin lab study on Pect (PMID: 30737130), we did not find a significant change in the overall PC and PE levels.

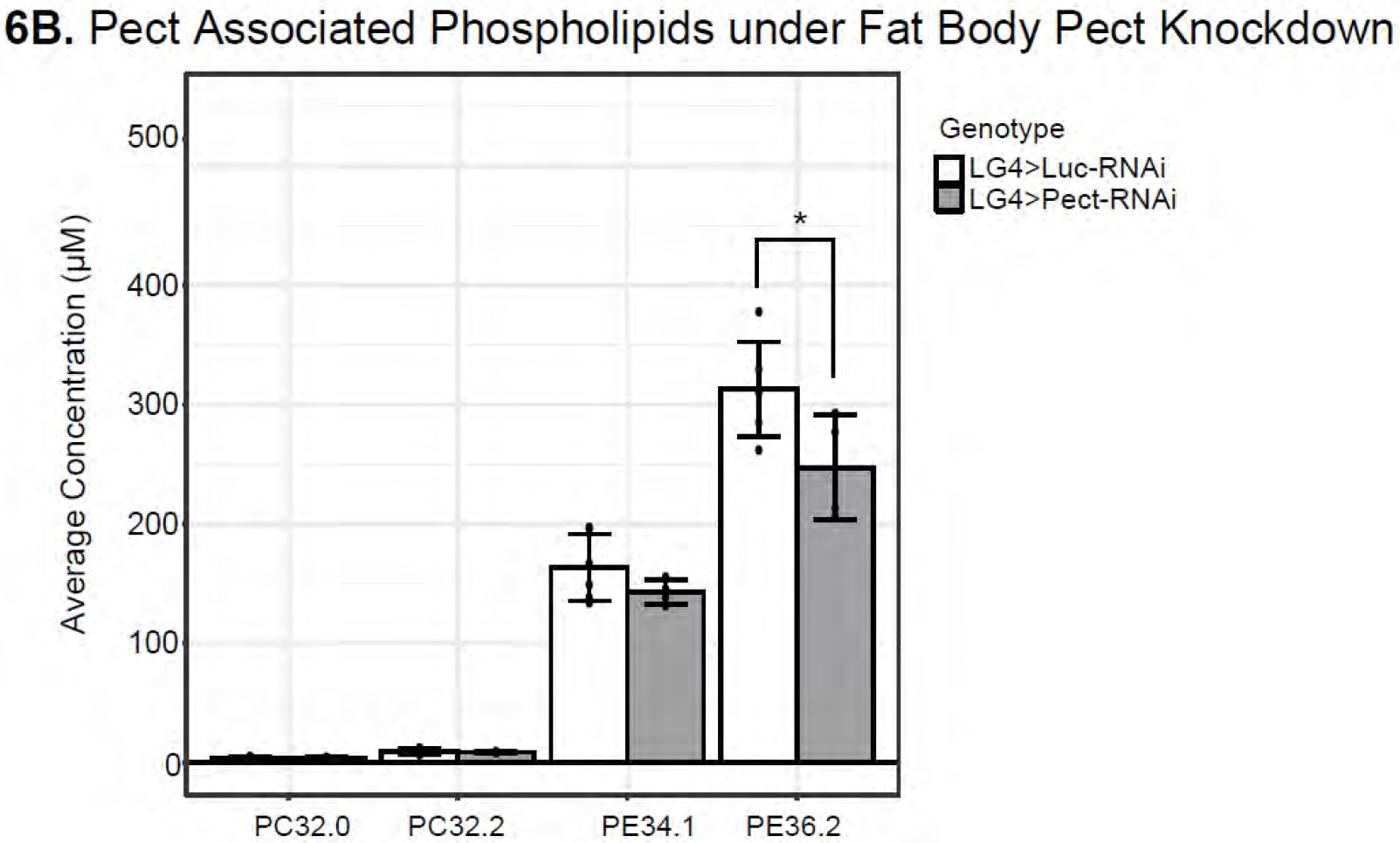
ii. Pect-KD shows increase in the phospholipid classes LPC and LPE (Figure 6A). In contrast, LPE is significantly downregulated on HSD Day 14 (Figure 3).

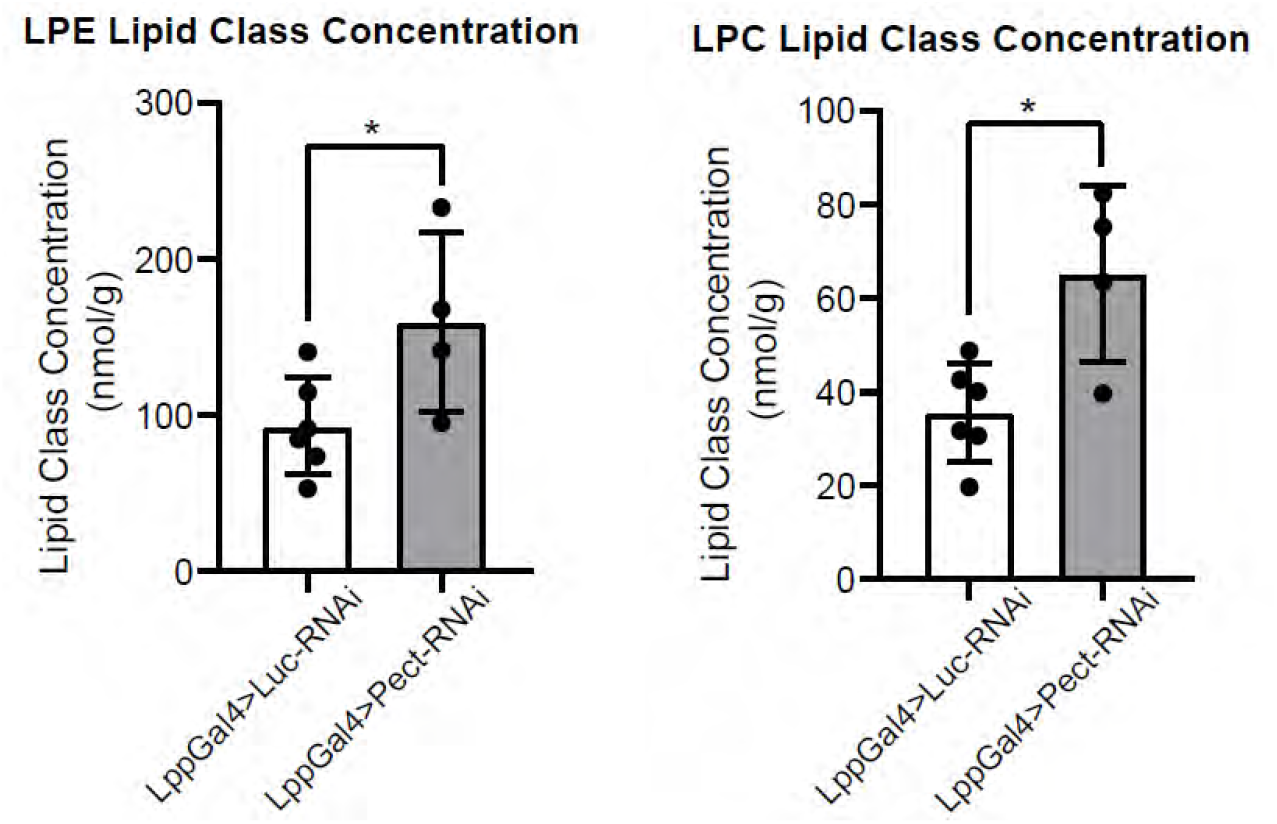

On 14-day HSD consistent with extreme upregulation of Pect mRNA fed flies (Figure S6A; Pect mRNA 200-250 fold), PE trends upwards on 14-day HSD (Figure 3) and PE 36.2 trends higher (Figure S7C). We note that on the surface of it PE and LPE *per se* are contrasting between 14-day HSD lipidome and fat-specifc Pect-KD. But there is a significant commonality that under both states there is an imbalance of phospholipids classes PE and LPE. Hence, we propose that maintaining the compositional balance of phospholipid classes PE and LPE is critical to hunger-driven feeding and insulin sensitivity. Hence, either increase or decrease, of these key phospholipid species, may lead to abnormal hunger-driven feeding.

Finally, fat-specific Pect-OE did not cause significant changes to lipid species (Figure S9). This could either be due to the fact that in fat-specific Pect-OE flies under normal food and that we were assaying whole body lipid levels and not fat-specific lipid changes. But to counter that, even a 60% reduction in Pect mRNA levels (Figure S6A), was sufficient to produce an effect on whole body phospholipid balance (Figure 6). Hence, we speculate that by maintaining a basally higher (7-fold higher Pect mRNA level Figure S6A), might allow 14-day HSD-fed flies to buffer the negative effects of HSD and we predict that it might take longer to disrupt the phospholipid balance and HDF response.

We have now included a section in the discussion - Page 14 Lines 26-34-under the subtitle “*The implications of relationship between Pect levels and HSD*”. We have pasted an excerpt from that subsection below for this reviewers assessment.

> *“Also, we note that over-expression of Pect cDNA in the fat-body does not alter phospholipid balance (Figure S9) and indeed improves HDF on HSD (Figure 7B). While this may appear inconsistent, it is critical to note that over-expression of Pect cDNA using UAS/Gal4 only increases Pect mRNA expression by 7-fold (Figure S6A), whereas HSD causes its upregulation by 250-fold (Figure S6B). Hence, we speculate that an increased ‘basal’ level of Pect such as by that provided by a cDNA over-expression in fat, may be protective to the negative effects of HSD (Figure 7B) without affecting overall phospholipid levels (Figure S9), but extreme upregulation Pect on HSD affects the PE and LPE balance (Figure 3).”*

2. A central hypothesis in the study is that the HSD over a period of 14 days results in insulin resistant and that these changes are leading to changes in hunger dependent feeding. I would encourage the authors to determine if Foxo mutants are resistant to these HSD-induced effects on HFD.

We thank the reviewers for this suggestion. However, given that dFOXO nuclear localization rather than expression levels regulate insulin sensitivity, we feel that disrupting dFOXO levels via mutation or knockdown will produce a plethora of indirect effects including developmental abnormalities (PMID: 24778227, PMID: 16179433, PMID: 29180716, PMID: 12893776). Our data suggest that chronic HSD treatment and Pect affect insulin sensitivity in fat tissue. However, we feel that investigating whether insulin sensitivity/FOXO signaling in fat tissue regulates feeding behavior is outside the scope of our work.

3. In lines 25-30, the authors draw the conclusion that an increase in unsaturated fatty acid species is associated with the HSD and that these changes results in a more fluid lipid environment. While I agree with the model, the manuscript contains no evidence to support such a model. Either test the hypothesis or move the last line of the section to the discussion.

We thank the reviewer for this important and insightful comment. We agree that the data we presented and discussed in the original version is at the moment speculative. Addressing the hypothesis that increase in unsaturated fatty acid species result in a more fluid lipid environment will require us to build tools and expertise. Hence, this hypothesis is better suited for exploration in a future study. Given this, we have moved this out of the results section into the Discussion section titled “*HSD and fat-specific PECT-KD causes changes to phospholipid profile*” (See excerpt below from page 13, lines 24-35).

> *“In addition to changes in phospholipid classes, we found that HSD caused an increase in the concentration of PE and PC species with double bonds (Figure S4C and S4D). Double bonds create kinks in the lipid bilayer, leading to increased lipid membrane fluidity which impacts vesicle budding, endocytosis, and molecular transport*^14,92^*. Hence it is possible that a mechanism by which HSD induces changes to signaling is by altering the membrane biophysical properties, such as by increased fluidity, which would have a significant impact on numerous biological processes including synaptic firing and inter-organ vesicle transport.”*

Also, as per the reviewer’s guidance, given that we are speculating here, we have also shifted this dataset from Main figure 4 to supplement S4C and S4D.

In addition, lines 25-30 state that FFAs are increased after 14 days of a HSD. Figure 3A shows the exact opposite - FFAs are significantly decreased in 14 day fed animals despite being elevated in the 7 day fed animals. This is an interesting result that warrants discussion. Moreover, I would encourage to examine the lipidomic data more carefully to ensure that the text accurately portrays the lipid profiles.

We apologize for misstating that FFAs are decreased on 14-day HSD in the lines 25-30. It was an error and we have corrected this. We agree with the reviewer that the reduction of FFA on Day 14-HSD is an intriguing and unexpected observation that needs to be emphasized and further discussed. To this end, we have added figure S4B, wherein we have provided the difference in FFA concentration (by species) after days 7 and 14.

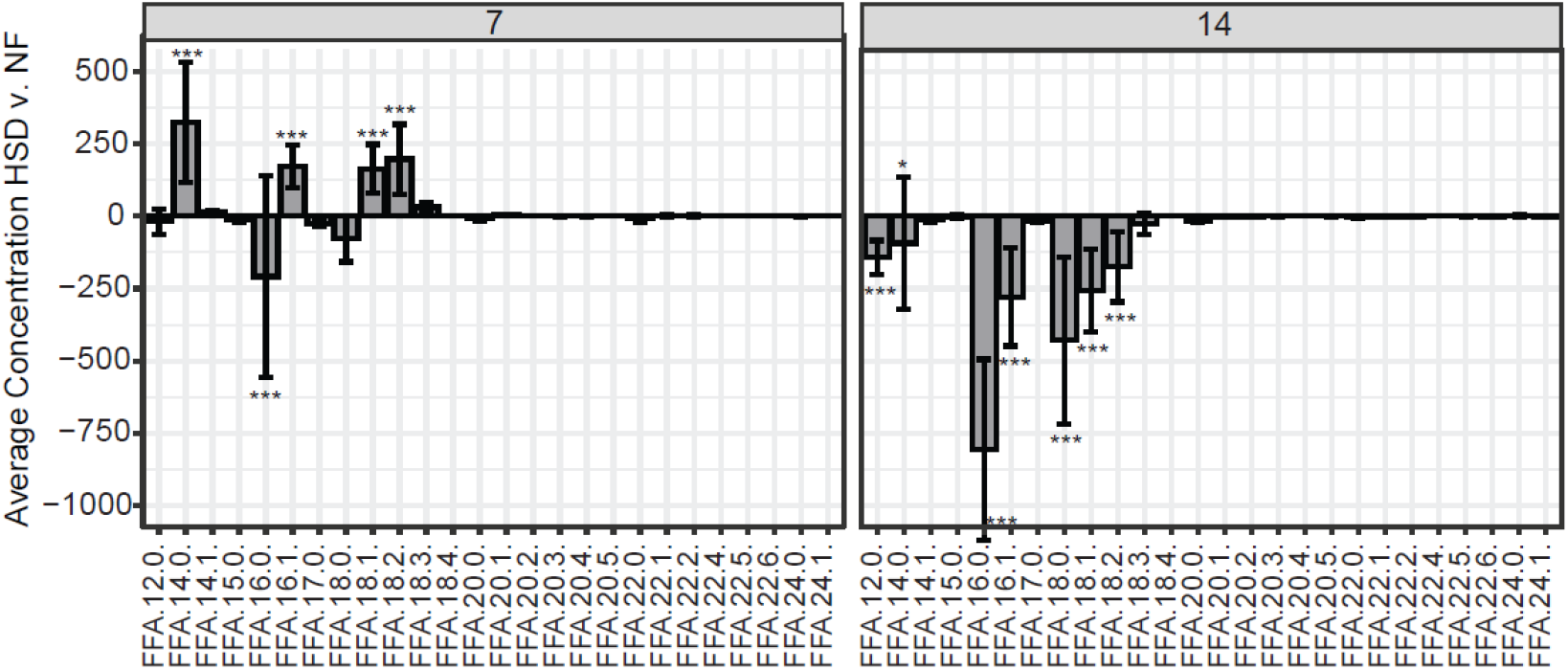

Furthermore, we have discussed what the potential meaning of reduced FFA at Day 14 implies in page 12, lines 19-27 of the Discussion section titled “*HSD and fat-specific PECT-KD causes changes to phospholipid profile*”. We have stated the following-

> “*We speculate that this reduction in FFA maybe due to their involvement in TAG biogenesis (PMID: 13843753). We were interested to see if the decrease in FFA correlated to a particular lipid species, as PE and PC are made from DAGs with specific fatty acid chains. However, further analysis of FFAs at the species level did not reveal any distinct patterns. The majority of FFA chains decreased in HSD, including 12.0, 16.0, 16.1, 18.0, 18.1, and 18.2 (Figure S4B). This data was more suggestive of a global decrease in FFA, likely being converted to TAG and DAG, rather than a specific fatty acid chain being depleted.”*

The processed lipidomics data should also be included as supplementary data table so that they can be independently analyzed by the reader.

We thank the reviewer for this suggestion. As per the reviewers request, we have included the raw data as an attachment in our supplementary material (Supplementary Files 1-3.), so that interested readers can use the datasets generated in this study for future work and further analysis.

Beyond these experimental suggestions, the manuscript needs significant editing for clarity. While I won’t provide a comprehensive list, the authors need to provide accurate descriptions and annotation of genotypes (including w[1118], which is written as W1118), typos, and formatting. I’ve listed a few examples below:

1. Page 3, Line 1 and 2: “…have been shown to impact feeding behavior and metabolism that leads to…” This is an awkward and grammatically incorrect sentence.
2. Page 3, Lines 7-32 is one very large paragraph but contains concepts that should be broken down over at least three paragraphs.
3. Page 3, Line 25: A description of the reaction catalyzed by Pect would be helpful for a manuscript focused on Pecte activity.
4. Page 4, Line 10: “previously characterized method of eliciting diet induced feeding behavior.” As stated in the text, the method is previously described yet the manuscript characterizing the method isn’t cited.
5. Figure legend 3 contains a random assortment of capitalized lipid species. Also, the names of lipid species are inappropriately broken into multiple names. Please use correct nomenclature throughout the manuscript.

The list above is nowhere near comprehensive. The manuscript requires significant editing.

We are grateful to the reviewer for drawing our attention to these errors. We have made significant edits to the revised manuscript to address the above-mentioned concerns, as well as made additional textual changes throughout and copyedited it. We hope that the reviewer will find the manuscript reads better and the clarity and preciseness is significantly improved.

Reviewer #2 (Significance (Required)):

I find the study to be potentially very important - the authors combine a longitudinal study that would be difficult in any other model with the powerful genetic tools available in the fly. The findings will significantly advance our understanding of how lipid metabolism links dietary nutrition with feeding behavior.

Once again, we are grateful for this reviewer’s thoughtful critique and encouraging words regarding our work and its potential impact.

**<u>Reviewer #3 (Evidence, reproducibility and clarity (Required)):</u>**

Summary:

This manuscript uses Drosophila to investigate how diet-induced obesity and the changes in the lipid metabolism of the fat boy modulate hunger-driven feeding (HDF) response. The authors first demonstrate that chronic exposure (14 days) of high sugar diet (HSD) suppresses HDF response. Through lipidome analysis, the authors identify a specific class of lipids to be elevated upon chronic HSD feeding. This coincided with the changes in expression of Pect, an enzyme that regulates the biosynthesis of these lipids. Modulating the expression of Pect specifically in the fat body affected HDF response.

We thank this reviewer for their rigorous and thoughtful critique and for identifying a key issue with our original study pertaining to a gap in how Pect mRNA levels on 14-day HSD are elevated but the Pect-KD phenocopies the HDF. Now by performing whole-body adult fly lipidomic on fat-specific Pect-KD we have resolved this issue and provided clarity on role of Pect in maintaining phospholipid homeostasis and thus subsequently impacts hunger-driven feeding. We hope the reviewer finds that the revised manuscript provides further clarity to the functional link between Pect’s role in fat-body and hunger-driven feeding.

Major comments:

The author claim that the HDF response in HSD is distinct between early (5d, 7d) and chronic (day 14) HSD feeding. However, the data seem to indicate that HDF response is significantly decreased at all time points in HSD. For example, at day 5 HDF response was increased only 3-fold in HSD (Figure 1C) compared to around 50-fold increase in NF (Figure 1B). The scale of the Y-axis in Figure 1B and 1C is an order of magnitude different. Including the starved data (NFstv and HSDstv) in Figure S1, normalized to NF fed group, would better visualize the overall trends. Related to this, having the source data for the actual number of feeding events would be useful (e.g., to see the baseline changes in feeding in different time points in Figure 1 and the effect of genetic manipulations in Figure 7).

As per the reviewers request, we now have modified our graphs to show source data (Figure S1) and show the raw feeding events.

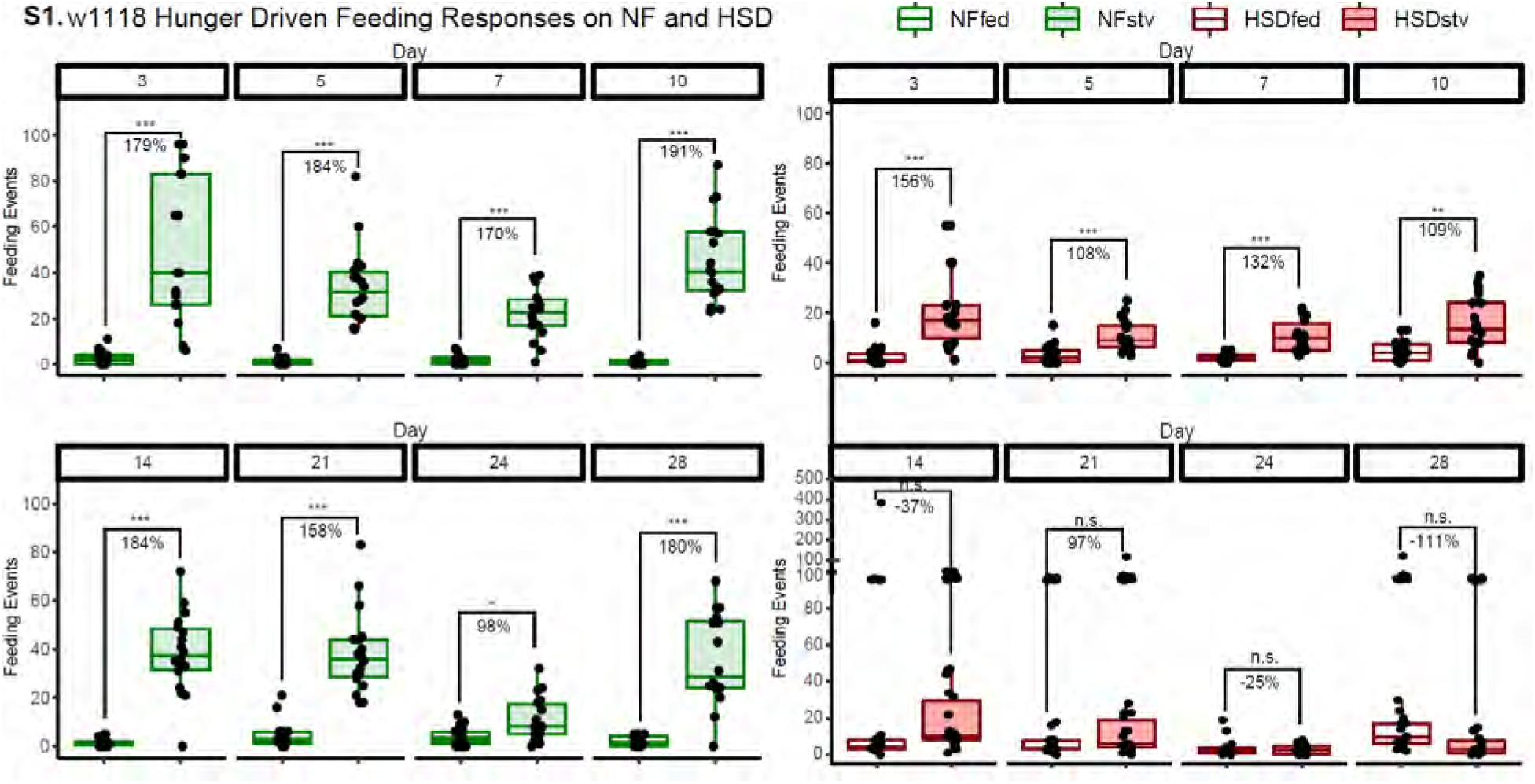

Then in the non-normalized graphs we plot, over a longitudinal time course, baseline and hunger-driven feeding events (Figure 1B-D). We also show that HSD fed flies do not display increased baseline feeding (Figure 1D) suggesting that the effect we see on HDF are no clouded by increased baseline feeding.

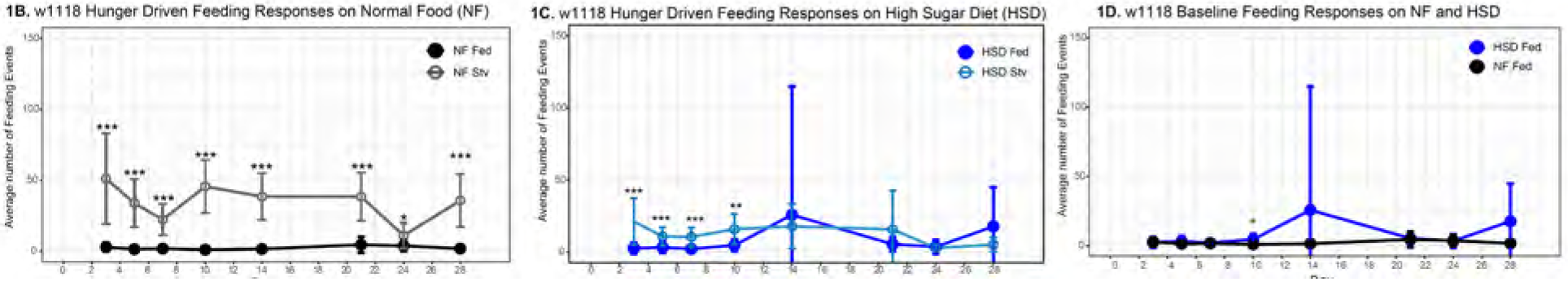

Yes, the reviewer makes an important point that HDF response on HSD fed flies is of a lower magnitude than NF fed flies. We think that is a biologically meaningful observation, as it suggests that flies have a remarkably fine-tuned ability to coordinate food-intake with nutrient store levels.

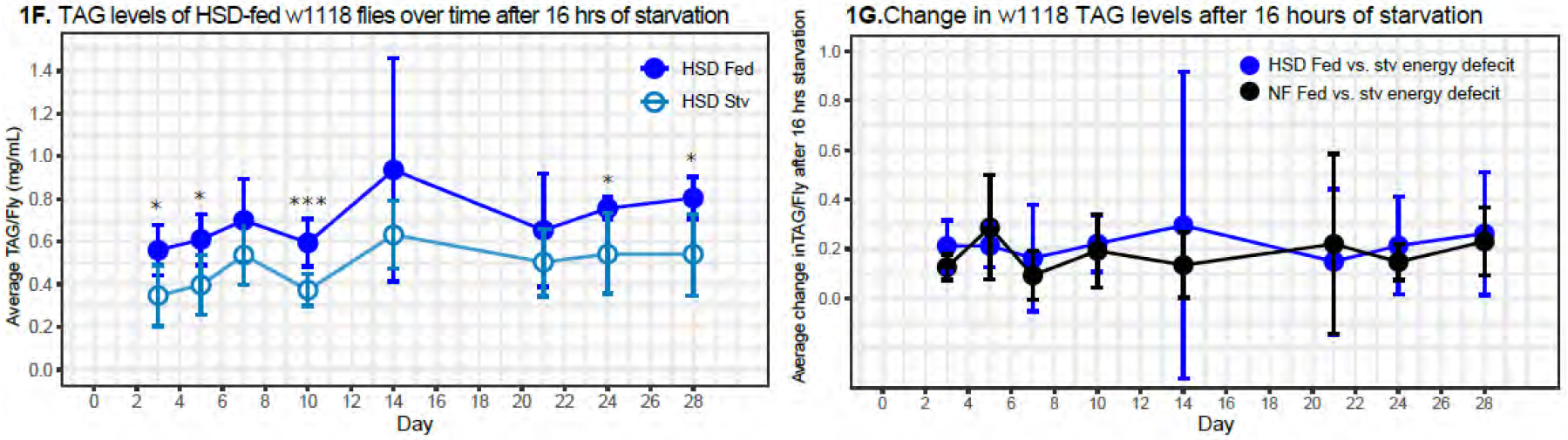

Now we have included a paragraph in the Discussion, Page 11 Lines 23-27, that say the following to ensure the readers appreciate this salient point raised by this reviewer.

> *It is to be noted that the HDF response of HSD-fed flies (Figure 1C, Days 3-10) is of lower order of magnitude than the NF-fed flies. This suggests that that in addition to sensing an energy deficit and mobilizing fat stores (Figure 1F, 1G, S1), HSD fed flies calibrate their starvation-induced feeding to compensate only for the lost amount of fat. Overall, this suggests that flies have a remarkably fine-tuned ability to coordinate food-intake with nutrient store levels*.

The association between fat body Pect level and phospholipid levels is not clear. Day 14 of HSD feeding shows high expression of Pect in the fat body and elevated levels of PC32.0 and PC32.2. The authors assume the high expression of Pect in the fat body is due to the compensatory response, but there are no data indicating downregulation of Pect levels at the earlier time points of HSD feeding. A previous study demonstrated that Pect mutant flies have lower levels of PC32.0 but higher PC32.2 (PMID: 30737130).

We agree that one puzzling aspect of the original version of this study was that Pect mRNA levels being very high on Day 14 HSD, but nonetheless the effects of Pect-KD phenocopied HSD. To resolve this, prompted by Reviewer #2 and #3 concerns, for this revised version we have now performed lipidomic analyses on whole adult flies, when Pect is knocked down (KD) by RNAi in the fat tissue. We now present a new dataset in Figure 6. Two striking changes occu. They are:

i. Pect-KD shows increase in the phospholipid classes LPC and LPE (Figure 6A). In contrast, LPE is significantly downregulated on HSD Day 14 (Figure 3).
ii. Pect-KD shows a significant reduction in specific class of PE 36.2 (Figure 6B). Our data regarding increase in PE 36.2 agree with a previous lipidomic analyses of Pect mutant retina (PMID: 30737130). In contrast, PE 36.2 trends upwards on 14 day HSD (Figure S7C) though not significantly.

On 14-day HSD consistent with extreme upregulation of Pect mRNA fed flies (Figure S6A; Pect mRNA 200-250 fold), PE trends upwards on 14-day HSD (Figure 3) and PE 36.2 trends higher (Figure S7C). We note that on the surface of it PE and LPE *per se* are contrasting between 14-day HSD lipidome and fat-specifc Pect-KD. But there is a significant commonality that under both states there is an imbalance of phospholipids classes PE and LPE. Hence, we propose that maintaining the compositional balance of phospholipid classes PE and LPE is critical to hunger-driven feeding and insulin sensitivity. Hence, either increase or decrease, of these key phospholipid species, may lead to abnormal hunger-driven feeding.

On day 14, HDF response was increased 70-fold in w1118 flies in NF (Figure 1B; w1118), but only 2.5-fold in lpp>LucRNAi control flies in NF (Figure 7A). This suggests that lpp-gal4 driver lines have a significant effect on HDF response. Using a different fat-body specific Gal4 line would be necessary to validate conclusions.

Regards reduced HDF magnitude, in our experience using UAS-Gal4 reduces HDF response magnitude consistently and cannot be compared to w1118 which is more robust. To account for background differences, we use Uas-Gal4 with control RNAi. It clearly shows differences in HDF response on starvation, but Pect and Pisd RNAi does not (Figure 7A). Hence, given that this experiment internally controls for any changes in HDF response for UAS-Gal4>RNAi, we conclude that HDF response in disrupted in Pect and PISD KD (Figure 7).

We only presented the Lpp-driver in our study, as this driver is the only fat-specific driver that has no leaky expression in other tissues, and is specific to fat as apolpp promoter used to generate this Gal4 line is only expressed in fat tissue (Eaton and colleagues, PMID: 22844248). Other widely used fat-specific drivers, including the pumpless-Gal4 (*ppl-Gal4*) driver has leaky expression in gut or other tissues (See Table 2 of this detailed study by Dr. Drummond-Barbosa https://www.ncbi.nlm.nih.gov/pmc/articles/PMC7642949/). If the reviewer is aware of a fat-specific Gal4 line, other than Lpp-Gal4, which has a highly specific expression in the fat tissue without leaky expression in other tissues, then we are happy to take onboard the reviewer’s suggestion and try that fat-specific Gal4 that they suggest.

HSD feeding promotes Pect expression (Figure S3C) and global changes in phospholipid levels (Figure 3, 4). Therefore, shouldn’t Pect overexpression (not Pect RNAi) in a normal diet mimic HSD feeding state and promote loss of HDF response? Conversely shouldn’t knockdown of Pect in HSD rescue loss of HDF response?

We agree that a puzzling aspect is that Pect mRNA levels are significantly elevated in HSD Day-14, but Pect-KD showed displays the inappropriate HDF response. As we have described in our response to this reviewer on Page 19, we believe that Pect-KD and HSD disrupt PE and LPE balance overall but in different ways. Whereas Pect-OE using cDNA expression in fat body does not cause a significant change to any lipid class (Figure S9), and our results suggest that basally higher level of PECT is likely to be protective on HSD with respect to HDF(Figure 7B).

To ensure that we appropriately discuss and clarify this issue, we have now included a section in the discussion - Page 14 Lines 26-33-under the subtitle “*The implications of relationship between Pect levels and HSD*”. We have pasted an excerpt from that subsection below for this reviewers assessment.

> *“Also, we note that over-expression of Pect cDNA in the fat-body does not alter phospholipid balance (Figure S9) and indeed improves HDF on HSD (Figure 7B). While this may appear inconsistent, it is critical to note that over-expression of Pect cDNA using UAS/Gal4 only increases Pect mRNA expression by 7-fold (Figure S6A), whereas HSD causes its upregulation by 250-fold (Figure S6B). Hence, we speculate that an increased ‘basal’ level of Pect such as by that provided by a cDNA over-expression in fat, may be protective to the negative effects of HSD (Figure 7B) without affecting overall phospholipid levels (Figure S9), but extreme upregulation Pect on HSD affects the PE and LPE balance (Figure 3).”*

We would have liked to test Pect protein expression on HSD, but since we were unable to access antibodies for Pect published in a prior study (PMID: 33064773) from Dr. Wang’s lab (see Page 10-11, of response to Reviewer #1). Hence, we were unable to test how the proteins levels of Pect correlate with the 250-fold increase mRNA expression.

In conclusion, we hope the reviewer appreciates that our results regarding Pect function are consistent with the main conclusion that achieving the right phospholipid balance between PE and LPE, is critical for an organism to display an appropriate HDF response.

Minor comments:

All graphs should plot individual data points and showed as box and whisker plot as much as possible.

Thanks for this suggestion, we have added individual data points to the vast majority of figures in the paper. We have made exceptions to graphs such as seen in figure 1 and FigureS4B-D where we find individual data points add an unnecessary layer of complexity. We hope these changes provide additional clarity and strength to the claims made in this manuscript.

Data for day 14 missing in Figure S4A and S4B.

We have provided Day 14 for the PC composition and PE composition, due to changes in Figures, they are now S7A and S7B.

Reviewer #3 (Significance (Required)):

The interactions between diet-induced obesity, peripheral tissue homeostasis and feeding behavior is an interesting topic that can be addressed using Drosophila. This manuscript demonstrates how fat body Pect levels affect HSD induced changes in hunger-driven feeding response. However, at this point, the functional association between fat body Pect level, global phospholipid level, and loss of hunger-driven feeding response in chronic HSD feeding is not clear.

We hope the revised data, and discussion of the paper, provides well-substantiated functional association on the importance of maintaining phospholipid balance, driven by Pect enzyme, as a critical regulator of hunger-driven feeding behavior. As stated in the revised discussion, the key take home message of our manuscript is that on prolonged HSD exposure PC, PE and LPE levels are dysregulated, the loss of phospholipid homeostasis coincided with a loss of hunger-driven feeding. Following this lead on phospholipid imbalance, we then uncovered a critical requirement for the activity of the rate-limiting PE enzyme PECT within the fat tissue in controlling hunger-driven feeding.

